# Separate and shared low-dimensional neural architectures for error-based and reinforcement motor learning

**DOI:** 10.1101/2022.08.16.504134

**Authors:** Corson N. Areshenkoff, Anouk de Brouwer, Daniel J. Gale, Joseph Y. Nashed, Jason P. Gallivan

## Abstract

Motor learning is supported by multiple systems adapted to processing different forms of sensory information (e.g., reward versus error feedback), and by higher-order systems supporting strategic processes. Yet, the extent to which these systems recruit shared versus separate neural pathways is poorly understood. To elucidate these pathways, we separately studied error-based (EL) and reinforcement-based (RL) motor learning in two functional MRI experiments in the same human subjects. We find that EL and RL occupy opposite ends of neural axis broadly separating cerebellar and striatal connectivity, respectively, with somatomotor cortex, and that alignment of this axis to each task is related to performance. Further, we identify a separate neural axis that is associated with strategy use during EL, and show that the expression of this same axis during RL predicts better performance. Together, these results offer a macroscale view of the common versus distinct neural architectures supporting different learning systems.

---

The central nervous system’s ability to learn new mappings between motor commands and sensory outcomes is critical for a majority of behaviours that we perform on a daily basis. Indeed, such learning is not only essential when developing new motor skills or refining existing ones (e.g. becoming a better piano player), but also when adapting to changes in the body or environment, or interacting with new objects. Given the diverse repertoire of sensory signals that can be used to drive sensorimotor learning, it is not surprising that this flexibility is supported by multiple distinct learning processes tuned to different forms of sensory feedback (Doya, 2000; Wolpert et al., 2011).

Two of the main processes known to drive sensorimotor learning are Error-based Learning (EL) and Reinforcement Learning (RL). These can be differentiated by the nature of the information that the brain uses as the “teaching” signal. In the case of EL, this teaching signal is a sensory prediction error encoding the directional difference between the actual versus expected sensory outcome of a motor command (Wolpert et al., 2011). This is the type of learning that drives improved performance in dart-throwing, for example, and is far-and-away the best studied form of motor learning. In the case of RL, the teaching signal is a reward prediction error encoding the difference between the actual versus expected value of the outcome of an action (Sutton and Barto, 2018; Lee et al., 2012; Wolpert et al., 2011). Often, when perfecting a movement, there is no directional error signal indicating precisely how the movement should be altered in order to achieve an intended outcome, and so the motor system must learn (assign credit to) which component of the action led to the success or failure of the movement. For example, when first learning to swing on a playground set, a child may explore different actions (e.g., wiggle their body, extend their legs, etc.) in an attempt to get the swing moving (success). If they are unsuccessful, there is no error signal indicating exactly how they should modify their actions, and the motor system must learn through reinforcement which component of the action is responsible for the failure.

Consistent with the notion that separable mechanisms govern EL and RL, both computational and neural work has suggested a neuroanatomical division of labour between the cortico-subcortical circuits that support these two main learning processes (Doya, 2000). In the case of EL, there is broad consensus that the cerebellum plays a critical role in the computation of sensory prediction errors used to drive sensorimotor adaptation (Wolpert et al., 2011) and that EL primarily results from implicit learning mechanisms mediated through cortico-cerebellar pathways. In the case of RL, it is well understood that the basal ganglia plays a critical role in the computation of reward prediction errors used to drive reward-based learning, and that RL primarily results from implicit learning mechanisms mediated through cortico-striatal pathways (Knowlton et al., 1996; Doya, 2000; Ramayya, 2014). However, it also recognized that these coarse neuroanatomical divisions do not wholly capture the diversity of processes that contribute to sensorimotor learning (Taylor and Ivry, 2014; Spampinato and Celnik, 2021). Recent behavioural studies have shown, for example, that cognitive processes such as strategy use, which are explicit and declarative in nature, can support both EL and RL (de Brouwer et al., 2018; Holland et al., 2018; de Brouwer et al., 2021), and that the use of such strategies can lead to better learning performance (de Brouwer et al., 2018, 2021). Together, this work suggests that both EL and RL are driven by at least two separate components to learning: (1) An implicit component that may be cerebellar-dependent (EL) or striatal-dependent (RL) in nature, and (2) An explicit, strategic component that is likely supported by task-general executive processes in higher-order association cortex (e.g., prefrontal cortex; Taylor et al., 2014).

Disentangling the explicit versus implicit contributions to learning, and how these components map onto the neuroanatomy supporting EL and RL, has been challenging for several reasons: First, many neural studies of motor learning, particularly work in animals, fail to distinguish between explicit and implicit learning processes, their underlying neural circuitry, and their relative contributions to performance (Spampinato and Celnik, 2021). Second, to date, neural comparisons between EL and RL have often been indirect, guided by sampling biases in neural recording sites across different studies and species, as well as the use of very different kinds of tasks to probe these two main forms of learning. Here, we directly compared learning-related changes in neural activity in two motor learning experiments in the same participants: An EL experiment that involved error-based visuomotor adaptation during reaching, and an RL experiment that involved the reward-based shaping of reach trajectories. Using supervised dimension reduction techniques on human functional MRI data and joint-embedding both tasks in a common neural space, we first isolate a connectivity dimension, specific to the learning process, that distinguishes activity associated with EL versus RL. Secondly, we identify a connectivity dimension, derived from the EL task, that is linked to explicit strategy use, and then show that the expression of this same dimension during the RL task predicts better performance. Together, these neural results provide characterization of the distinct versus overlapping activity patterns associated with EL and RL, and are consistent with the view that explicit learning is supported through a common neural architecture.

## 1 Results

### 1.1 Behavioral tasks

Subjects (N = 36) performed an error-based learning (EL) task and a reinforcement learning (RL) task during two separate MRI testing sessions, one week apart. Both tasks required the subject to move a cursor, representing their finger position, to intercept a target using an MRI-compatible touchpad, subject to different forms of visual feedback.

In the EL task, subjects performed a classic visuomotor rotation task (Krakauer, 2009) wherein they performed center-out directed movements, using their right finger, towards a target that could appear in one of eight locations on a circular ring. After a baseline block (40 trials), subjects performed a learning block (160 trials) in which the correspondence between finger motion on the touchpad and movement of the cursor was rotated clockwise by 45°. Thus, subjects needed to learn to adapt their movements by applying a 45° counter-clockwise rotation in order to hit the target. At the conclusion of each trial, subjects received visual (error) feedback consisting of both the target location, as well as the position of their cursor on the target ring. Following the learning block, subjects performed 16 “report” trials (Taylor et al., 2014), in which they were asked to indicate their aim direction using a dial, controlled by their left hand, prior to executing their movement (via their right hand). These report trials were presented at the end of the task, after learning had taken place, in order to avoid drawing attention to the nature of the manipulation and thus biasing subjects’ learning behaviour (de Brouwer et al., 2018). Critically, these report trials provided us with an index of subjects’ explicit knowledge of, and strategy in counteracting, the visuomotor rotation (Taylor et al., 2014).

In the RL task, we used a modified version of a motor learning RL task (Wu et al., 2014) wherein subjects used their right finger on the touchpad to trace, *without visual feedback of their cursor*, a rightward-curved path (from a start position to target line). Following a baseline block (70 trials), in which subjects did not receive any feedback about their performance, they performed a learning block (200 trials) wherein they were instructed that they would receive score feedback, between 0 and 100, based on how accurately they traced the rightward-curved path. Critically, however, unbeknownst to the subjects, the score they received was *actually* based on how accurately they traced the mirror-image path (reflected across the vertical axis). Prior to beginning this learning block, subjects were instructed to maximize their scores across trials. As the cursor was invisible, subjects had only the score feedback (and thus no directional information) with which to refine their movements, and maximize their score, across trials. Illustrations of both tasks, as well as typical subject performance, are shown in Figure 1a-d.

**Figure 1:**
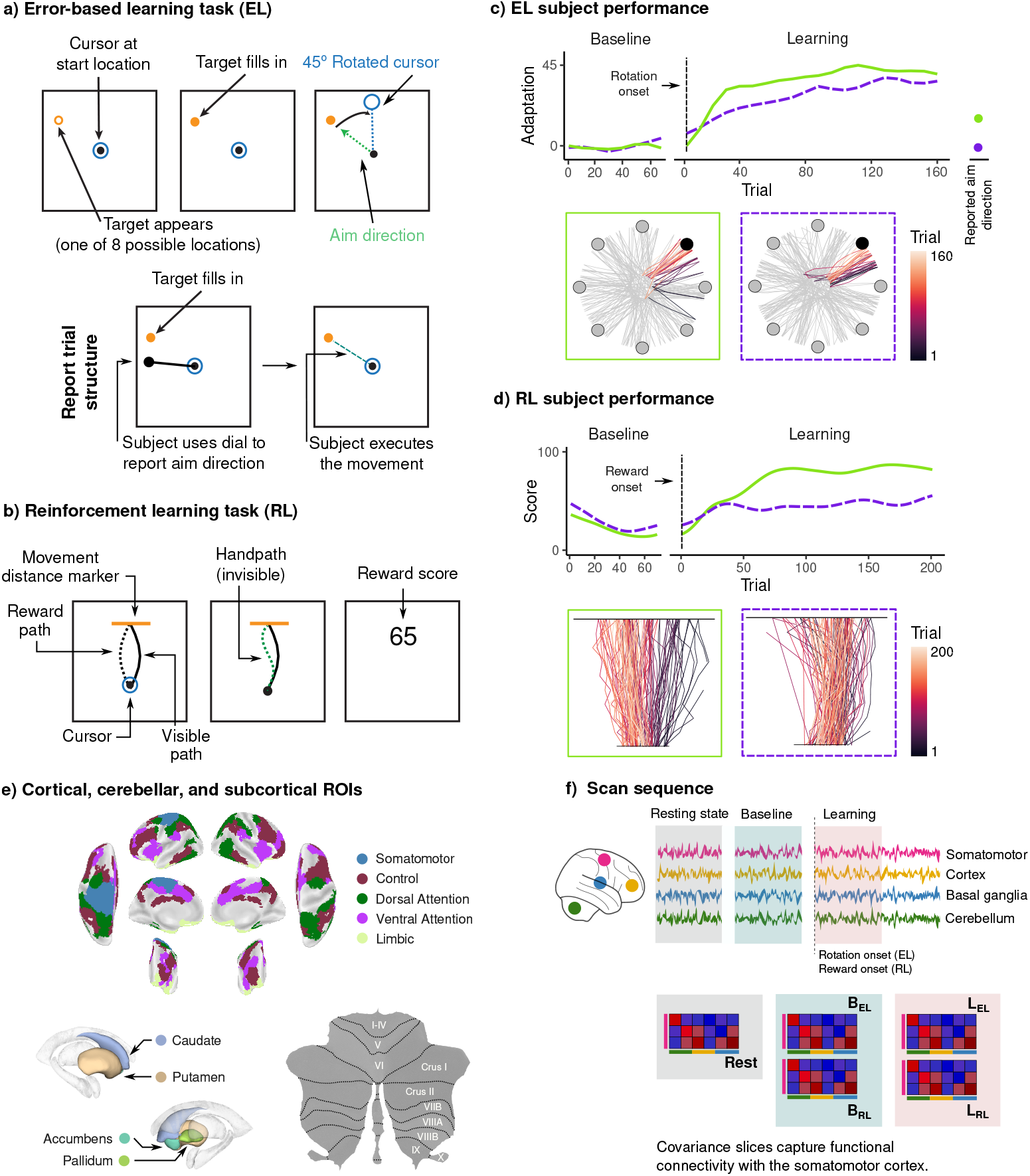
Experimental methods and data analysis. **a)** Trial structure for the error-based learning (EL) task. On each trial, subjects moved a cursor, representing their right finger position, from a start location to a target, using an MRI-compatible touchpad. (Top) During learning (rotation) trials, the mapping between finger movement and cursor direction was rotated by 45°. (Bottom) At the end of the task, subjects performed 16 ‘report’ trials in which they reported their intended aim direction using a dial, controlled by their left hand, before executing the movement. This provided us with a measure of subjects’ explicit strategy in counteracting the rotation. **b)** Trial structure for the reinforcement learning (RL) task. On each trial, subjects traced a curved (visible) path from a start location to a target line, without visual feedback of their cursor. After a baseline block of trials, subjects were then instructed that they would receive score feedback, presented at the end of the trial, based on their accuracy in tracing the visible path. However, unbeknownst to subjects, the score they received was *actually* based on how accurately they traced the mirror-image path, which was invisible to participants. **c)** Example subject data for a faster (in green) versus slower (in purple) learner in the EL task. (Top) Smoothed adaptation curves, along with each subject’s reported aim direction relative to the target (at far right). (Bottom) Cursor trajectories for individual trials. A subset of trials consisting of movement to the filled-in target have been colored by trial number to demonstrate that the slower learner (purple) tended to adapt more gradually over the course of the learning block. **d)** Example subject data for the faster (in green) and slower (in purple) learner in the RL task. (Top) Smoothed score curves. Note that reward scores were not presented to participants during the Baseline block. (Bottom) Cursor trajectories for individual trials (which was invisible to participants). Note that the better learner was able to shift their path towards the rewarded (mirror-image) path. **e)** For our neural analyses, we studied functional connectivity between regions located in the left somatomotor cortex, bilateral task-positive functional networks across the cortex, as well as regions in the striatum and cerebellum. **f)** For each region of interest (ROI), we estimated its covariance with every other regions in the brain during each of the resting state, baseline, and early learning epochs. We then extracted the slices of these networks corresponding to covariance between the somatomotor cortex and the remaining ROIs.

Note that all subjects performed the RL task during the first testing session (i.e., testing order was not counterbalanced). We made this decision in light of our prior evidence that subjects alter their behavior after the performance of explicit report trials in the EL task (de Brouwer et al., 2018), often developing explicit knowledge of the rotation (and thus enhancing their performance). We were concerned that, had subjects performed the EL task first, they may have anticipated a similar manipulation in the RL task, and thus would not have approached the task in a naive fashion. Also note that, prior to performing the RL task in the first session, subjects completed a single resting-state scan to allow for the measurement of each subjects’ intrinsic patterns of functional connectivity (see more on this below).

Behavioral data associated with the two motor learning tasks are shown in Figure 2. For each subject, we obtained learning trajectories by recording the adaptation on each trial of the EL task (defined as the hand direction relative to the target) and the score obtained on each trial of the RL task. Mean performance and individual subject learning curves are shown in Figure 2a, confirming that subjects, on average, successfully learned each task. However, these group-averaged results are potentially misleading, as they obscure significant inter-subject variability in both the rate and extent of learning in each task. To summarize subject-level performance in a data-driven manner, and isolate the dominant patterns of variability in each task (see Methods), we performed functional principal component analysis (fPCA; Shang, 2014) on subjects’ learning curves. These components, as well as subjects’ loading (scores) onto these components, are shown in Figure 2b. For each task, we found that a single component, encoding overall learning, captured the majority (75%) of variability in subjects’ learning curves. Thus, these learning performance scores formed the basis of our analyses moving forward.

**Figure 2:**
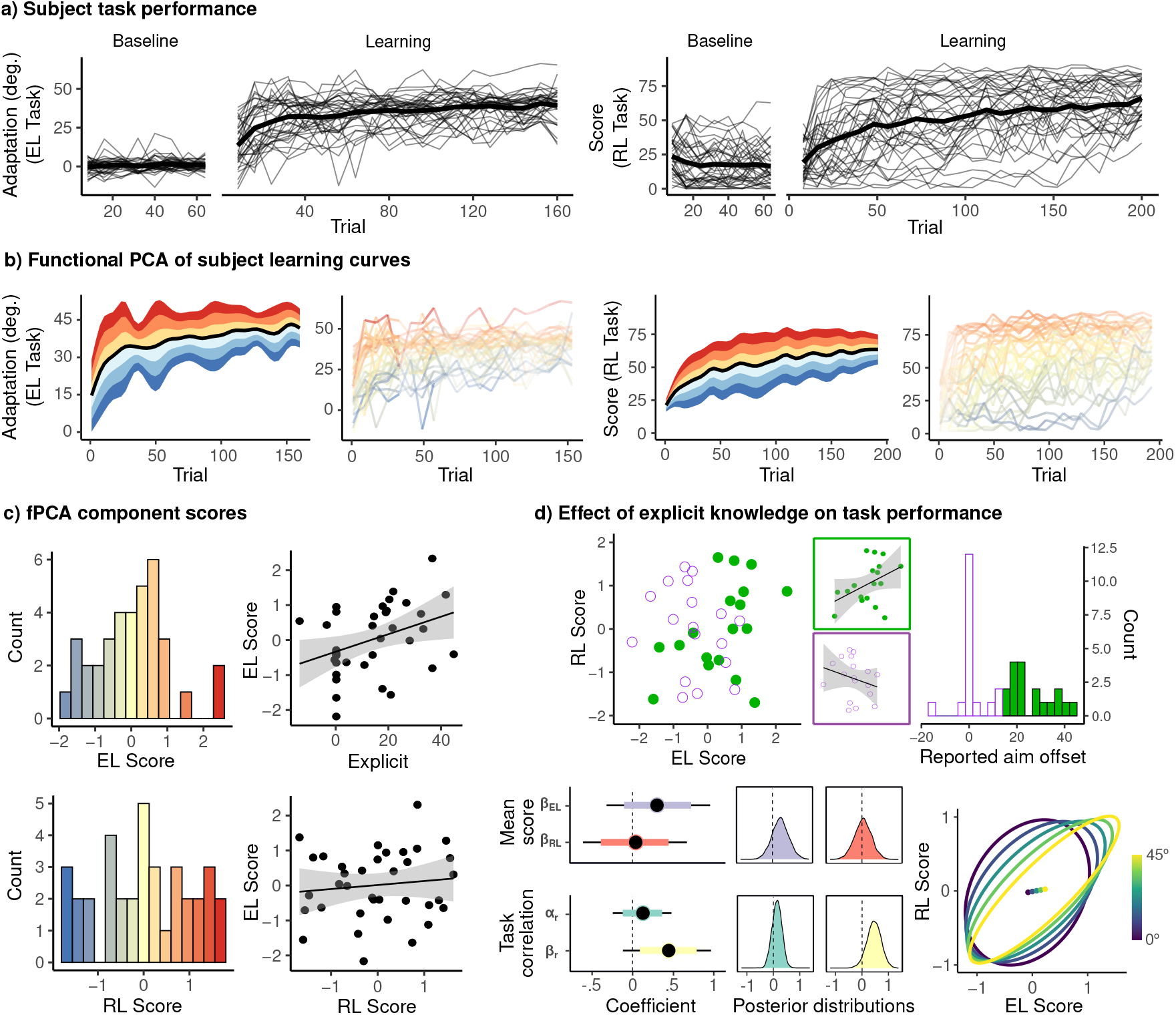
Subject learning behavior. **a)** Individual subject learning curves for the EL (left) and RL (right) tasks. Solid black lines denote means across all subjects. **b)** We performed functional principal component analysis on subjects’ learning curves in each task in order to identify the dominant patterns of variability. In both cases, a single component – encoding overall learning – explained a majority of the observed variance. Left panel shows the top component, with red (resp. blue) bands denoting the effect of positive (resp. negative) scores, relative to mean performance. In both tasks, the top component encoded overall task performance (better or worse than average). Right panels show individual subject learning curves, colored by subjects’ score on the top component. **c)** (left) Histograms of subjects’ scores on the EL and RL components. (right top) Subjects reporting explicit knowledge of the rotation exhibited greater performance in the EL task. (right bottom) Relationship between EL and RL task performance. **d)** (top) Relationship between EL and RL task performance after median split of subjects by their explicit reports. Among highly explicit subjects (purple), performance in EL and RL tasks was highly correlated. (bottom) We constructed a Bayesian multivariate regression model in which EL and RL performance, as well as the error covariance, varied as a function of subjects’ reported aim direction. Left panel shows posterior means (black dots) and highest posterior density intervals (thick and thin lines, 50% and 90% respectively). Top intervals are standardized coefficients for the effects of explicit knowledge on mean EL and RL performance. Bottom intervals are the intercept and slope for the effect of explicit knowledge on the (Fisher transformed) correlation. (middle panel) Posterior distributions for the parameters just described. (right panel) Posterior mean predicted distributions of EL and RL scores as a function of reported aim offset. Points denote predicted mean scores, while ellipses represent a Mahalanobis distance of 1.

In the EL task, we observed a large degree of variability in subjects’ reported aim directions during the report trials, ranging from zero explicit knowledge (wherein subjects reported aiming directly at the target), to overestimation of the aiming direction required to fully counteract the visuomotor rotation (greater than 45°). We acknowledge that this reporting procedure provides an imperfect measure of explicit strategy use, as subjects who report aiming directly at the target may still possess some explicit knowledge of the rotation, but may simply fail to understand what is being asked of them during the report trials. However, as these reports are known to correlate with several features of performance associated with the explicit component of learning (de Brouwer et al., 2018, 2021), they nonetheless provide us with a quantifiable index of subjects’ strategy use when performing the task.

Subject scores on the EL and RL performance components are shown in Figure 2c. Consistent with prior work (de Brouwer et al., 2021), we observed only a modest positive correlation between subjects’ performance in EL and RL (*r* = .11, *t*_34_, *p* = .5). However, further inspection reveals that this correlation may have been mediated by subjects’ use of cognitive strategies: Among subjects who reported using a re-aiming strategy during the EL task, we found that performance across the two tasks was highly correlated. Conversely, among subjects’ reporting no explicit knowledge of the rotation, we observed zero correlation between EL and RL performance. For a visualization of this effect, Figure 2d (top) shows the association between EL and RL performance based on a median split of subjects’ explicit reports, showing a clear positive relationship in highly explicit subjects, and no relationship at all for subjects reporting little explicit knowledge. To quantify this effect, we fit a Bayesian multivariate regression model in which both EL and RL task performance, as well as the correlation between them, varied as a function of subjects’ explicit reports (see Methods for details). We found evidence of a positive effect of explicit strategy use on the correlation between EL and RL task performance, as well as a moderate overall effect of explicit strategy use on EL task performance (Figure 2d). This result is consistent with prior evidence that explicit strategy use is associated with better learning performance across different tasks (de Brouwer et al., 2018, 2021).

The above behavioural findings suggest that subjects can flexibly recruit several distinct learning systems during EL and RL. Some of these systems, such as error-based or reinforcement learning, are specialized for handling different forms of feedback (sensory and reward prediction errors, respectively), and thus are expected to be selectively recruited during the performance of one task versus the other (i.e., task-specific). Alternatively, subjects may also recruit task-general, explicit strategic processes (Anguera et al., 2010; Taylor et al., 2014; McDougle and Taylor, 2019; Christou et al., 2016; Spampinato and Celnik, 2021) to bolster learning performance. Subjects who do not recruit such task-general systems during learning must (necessarily) rely on implicit, task-specific processes, and so may succeed or fail in one task independently of the other. By contrast, the recruitment of task-general, strategic processes during learning are expected to introduce a correlation in the performance across tasks.

Using these above distinctions as a guidepost, in the next sections we study learning-dependent changes in functional connectivity between the somatomotor cortex and a variety of regions implicated in implicit error-based adaptation, reinforcement learning, and higher-order cognitive processes. Our goals are twofold: First, we aim to characterize the unique activity patterns supporting learning in each task. Second, we aim to isolate activity patterns associated with explicit strategy use in the EL task. If such strategies are indeed supported by task-general executive processes, then the recruitment of these same activity patterns during RL should be associated with higher learning scores in the RL task.

### 1.2 Supervised embedding reveals EL- and RL-specific patterns of functional connectivity

As noted above, successful learning may involve not only the engagement of implicit error-based and reinforcement-based learning systems (our focus in this section), but also the recruitment of higher-order, task-general systems. These task-general systems, in turn, may not only include explicit learning processes (e.g., strategy use), but also general processes related to task-engagement, attention, effort, and the perceptual processing of task stimuli. Changes in task difficulty, associated with the onset of the visuomotor rotation (EL task) and the presentation of score feedback (RL task), may likewise be associated with neural changes that are not directly related to implicit or explicit learning processes per se. For this reason, isolating functional networks associated with EL and RL is more challenging than simply comparing functional connectivity between tasks, or comparing the baseline and learning phases in each task individually. Specifically, as the baseline phases of each task assay brain activity *prior* to learning - that is, they do not include score feedback (RL), nor involve subjects making large systematic directional errors (EL) - they should not differ in the relative engagement of RL and EL systems. Conversely, purely task-general activity should not distinguish between the two tasks at all, but should nevertheless distinguish between the resting-state and task-based scans, as well as between the baseline and learning task blocks. We thus used a supervised variant of principal component analysis (sPCA; Barshan et al., 2011) to search for low-dimensional neural structure that distinguishes only between the learning epochs of the EL and RL tasks (a task-unique connectivity axis), which required that we isolate this activity from low-dimensional neural structure that distinguishes between rest, baseline and the learning epochs (a task-positive connectivity axis). Because the latter axis conflates the explicit learning processes of interest with other task-general processes unrelated to learning (as mentioned above), our focus in this current section will be on isolating the task-unique axis. In the following section, we will turn our attention to isolating activity patterns selectively linked to explicit learning.

For our analyses, we focused on changes in the covariance between the somatomotor cortex and a network of cortical and subcortical regions implicated in both error-based (e.g., cerebellum) and reinforcement (e.g, striatum) learning, as well as higher-order cortical association regions. This was done so as to capture (1) top-down interactions between cognitive and somatomotor regions during learning, and (2) functional interactions between cortical somatomotor regions and subcortical areas commonly associated with EL and RL (the cerebellum and striatum). By constraining our analysis in this way, we sought to specifically characterize the degree to which each of the non-motor cortical regions (in cortical, cerebellar and striatal cortex) modulate their connectivity with areas located in the final motor output pathway (in somatomotor cortex) during task performance. To this end, we extracted cortical regions derived from the parcellation of Schaeffer et al. (Schaefer et al., 2018,; 400 region parcellation), as well as cerebellar and striatal regions extracted from the Diedrichsen (Diedrichsen, 2006) and Harvard-Oxford parcellations (Frazier et al., 2005; Makris et al., 2006), respectively. Within the cerebral cortex, we restricted our analysis to the contralateral (left) somatomotor cortex (Somatomotor A; 19 regions), and bilateral cortical regions belonging to task-positive cognitive networks, as defined by Yeo et al. (2011) (dorsal and ventral attention, control and limbic networks; 188 regions). This restricted set of ROIs (as opposed to a whole-brain analysis) had a dual purpose. First, it eased the interpretation of the results, as a full whole-brain analysis would include a great number of regions whose roles in the tasks are not clear, requiring either (1) a large number of comparisons in order to disentangle specific regions whose functional connectivity is meaningfully associated with each task, or (2) high-level summaries of distributed effects, which may not meaningfully speak to the roles of individual regions. Second, the high-dimensionality of a full whole-brain analysis would require both longer time windows in order to obtain precise covariance estimates, and it would make it more difficult to extract meaningful connectivity axes without a larger sample.

For each subject, we estimated covariance networks during the resting state, baseline, and learning scans in each task. As covariance estimates are biased, and this bias is a function of the sample size, we chose time windows of equal length (100 TRs) during each scan. During the learning blocks, we chose the beginning of the scan for our analyses given that: (1) this early learning period is typically associated with the most rapid improvement in performance (Taylor et al., 2014; de Brouwer et al., 2018, 2021), and (2) in the visuomotor rotation learning literature specifically, this early learning period is the point in time in which the contribution of explicit processes to learning is greatest (Taylor et al., 2014). Given that a large body of recent literature suggests that functional connectivity is dominated by static, subject-level differences that can obscure any task-related variance (Gratton et al., 2018), we first centered the covariance matrices (Zhao et al., 2018,; see Methods for details, and Figure 3a). Figure 3b shows the effects of this centering procedure. Note that, before centering, the covariance matrices exhibit strong subject-level clustering, which confounds any task-related structure. However, after centering, this subject-level clustering is abolished, and the task-related differences become more clear..

**Figure 3:**
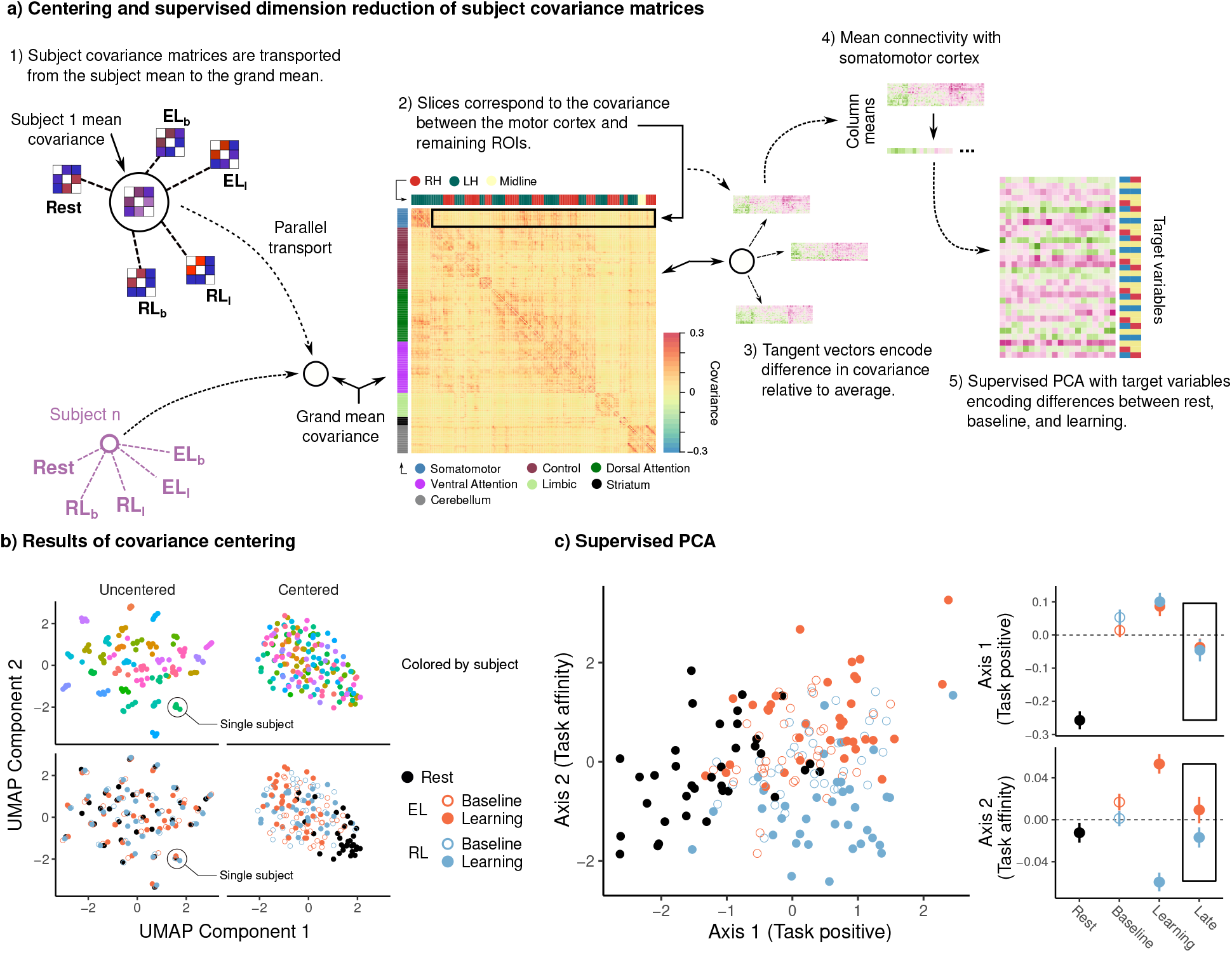
Supervised PCA reveals task-positive and task-specific patterns of functional connectivity with the somatomotor cortex. **a)** Covariance matrices estimated during rest, baseline, and learning were centered to align subject means to the grand mean covariance. We then projected the centered covariance matrices onto the tangent space at the grand mean in order to obtain tangent vectors encoding the change in covariance relative to the mean. We extracted slices of these tangent vectors containing the functional connectivity between the somatomotor cortex and the remaining regions (cortical, cerebellar and basal ganglia), and performed supervised PCA in order to isolate task-common and task-unique patterns of connectivity. **b)** Effects of covariance centering illustrated by UMAP. Each point denotes a single covariance matrix, colored by subject (top) or task epoch (bottom). Note the strong subject-level clustering before centering, which obscures any task-related effects. **c)** Results of supervised PCA. Each point denotes a covariance matrix. The first connectivity axis (which we call the *task positive* axis) increases monotonically from rest to baseline to learning. By contrast, the second connectivity axis *(task affinity)* distinguishes between learning in the EL and RL tasks. Note that projection of the covariance at late learning (the end of the learning scan) onto these axes revealed a general return to baseline-levels of expression for both axes (see bounded box at right).

Next, we projected each centered covariance matrix onto the tangent space around the grand mean covariance in order to obtain a tangent vector encoding the difference in functional connectivity relative to the grand mean. This approach has two advantages: First it isolates *differences* in covariance relative to the average; and second, while the space of covariance matrices is curved, the tangent space is flat, and thus operations such as principal component analysis, which assume that observations are constructed additively from the estimated components, are more sensible. Finally, we extracted a slice of the resulting tangent vector corresponding to the covariance between the somatomotor cortex and the remaining brain regions of interest (cortical, cerebellar and striatal regions). These slices were then vectorized and subjected to sPCA, according to the approach described above.

The scores on each connectivity axis for each subject and each scan are shown in Figure 3c, revealing that sPCA successfully extracted activity patterns satisfying our criteria. Specifically, note that the mean score of axis 1 increases monotonically from rest to baseline to learning, whereas the mean score of axis 2 shows a more minimal change from rest to baseline, and instead mainly distinguishes between the learning phases of the EL and RL tasks. Henceforth, we will refer to these connectivity dimensions as the *task positive* and *task affinity* axes, respectively.

Although we describe these connectivity axes above as reflecting activity patterns associated with *learning*, it is likewise possible that these axes, particularly the task affinity axis, mainly reflects the unique features of *motor execution* associated with each task. That is, if the task affinity axis simply encodes the differences associated with *performing* each task, then we would expect the expression of that axis to be similar during both the beginning (early) and end (late) of the learning epoch. In order to assess this, we separately projected the functional connectivity estimated during the end of the learning epoch in each task onto the two axes estimated from the rest, baseline, and early learning epochs. As can be seen in Figure 3c, we find that the expression of both axes generally returns to baseline-like levels by the end of the learning block. This suggests that the axes – in particular, the task-affinity axis – do not solely reflect differences in the *execution* of each task, but rather differences in neural structure associated with the early learning phase.

Surface maps for the task-positive axis are shown in Figure 4a, revealing the mean connectivity of each ROI with the somatomotor cortex. As can be seen in this figure, this axis is characterized by largely positive loadings across the cortex, striatum, and cerebellum (in particular, the anterior lobules; Nitschke et al., 1996; Stoodley and Schmahmann, 2010). Within the cerebellum, anterior regions showed increased connectivity with the somatomotor cortex during task, consistent with its established role in motor control. In contrast, limbic regions, comprising the temporal poles and oribitofrontal cortex, loaded negatively. Surface maps for individual subnetworks are included in Supplemental figures 1 and 2, respectively, for the task positive and task affinity axes.

**Figure 4:**
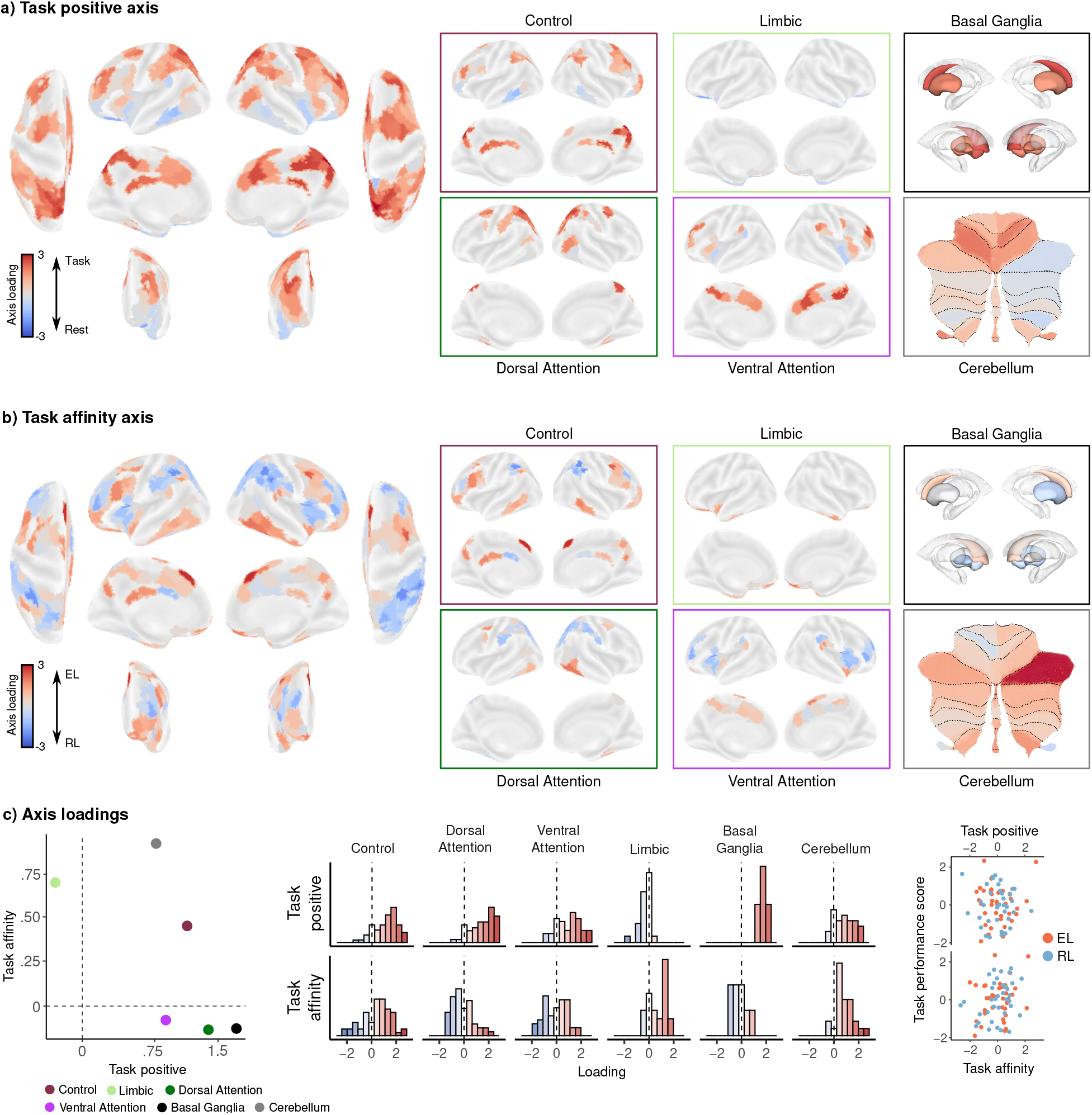
Surface maps for task-positive and task-affinity connectivity axes. **a/b)** Cortical, striatal, and cerebellar surface maps for the task-positive and task-affinity axes. **c)** (Left) Mean loadings on each axis for each network. (Middle) Histograms of individual ROI loadings for each network. (Right) For each task, we examined the relationship between learning performance (behavioral component scores) and the scores/loadings on the task-positive and task-affinity axes. We observed significant relationships between the task-affinity component and performance. Note that, for the RL task, the sign of the task-affinity score has been flipped so that the positive direction always denotes greater alignment with the respective task.

For the task-affinity axis, we describe ROIs with positive and negative loadings as being *EL-aligned* and *RL-aligned*, respectively; meaning that the connectivity of each ROI with somatomotor cortex is greater (resp. less), on average, during the EL (resp. RL) task (see Figure 4b for surface maps). Here, we found that the cerebellum was overwhelmingly EL-aligned, consistent with its well-established role in error-based learning (Wolpert et al., 2011). Moreover, we found that this EL-alignment was greatest in the cerebellar hemisphere ipsilateral to the response (right) hand, consistent with its connectivity to the contralateral motor cortex (Buckner et al., 2011). The alignment of the striatum, by contrast, was weaker and more mixed across the two tasks, but with its ventral regions (in particular, the accumbens) being RL-aligned, consistent with its role in reward processing. At the cortical level, we note that the frontal eye fields were largely EL-aligned, consistent with their role in the orienting of spatial attention and findings showing shifts in gaze orientation during visuomotor rotation learning (see de Brouwer et al., 2018, for a discussion of these processes in the context of the VMR task). Within the control and dorsal attention networks, we observed a clear division between frontal regions, which tended to be EL-aligned, and temporal and parietal regions, which were largely RL-aligned. Notably, the ventral attention network showed a medial-lateral division between EL and RL aligned regions, respectively. Somewhat unexpectedly, we found that the limbic network, including the temporal poles and orbitofrontal cortex, which are commonly associated with reward-related processing (Behrens/Rushworth reviews), were largely EL-aligned (see Figure 4c)

We next asked whether the degree to which subjects express these task-affinity and task-positive axes is associated with performance in each task. To the extent that successful learning requires orienting the brain’s functional architecture to the unique demands of each task, then we would expect performance in EL and RL to correlate with subjects’ expression of the task-affinity axis. In order to test this, for each axis, we fit a linear mixed-effects model predicting subjects’ EL and RL performance scores as a function of their axis score during the learning phase of the respective tasks (with random intercepts for each task). We found a significant effect of task affinity (*β_std_* = .23, *t*_70_ = 2.66, *p* = .047), but not of task-positivity (*β_std_* = .009, *t*_70_ = −.074, *p* = .94), suggesting that better performance in each task is associated with the recruitment of activity patterns specific to each task.

To summarize, our task-affinity axis revealed that learning in the EL task was associated with increased connectivity between the somatomotor cortex and the cerebellum, and frontal and temporal cortical regions within the control, limbic, and dorsal attention networks. Learning in the RL task, by contrast, was associated with increased connectivity between the somatomotor cortex and the ventral striatum, ventral attention network, and parietal regions of the control and dorsal attention networks. We also found that the expression of this task-affinity axis was associated with better performance in each task. In addition, our task-positive axis showed that the transition from rest to task was associated with a near-global increase in functional connectivity between the somatomotor cortex and cortical, anterior cerebellar, and striatal regions, but that the expression of this axis was not associated with learning performance. However, as we noted at the outset, our identification of the task-positive axis is likely to conflate explicit learning processes (of interest here) with task-general processes unrelated to learning (e.g., attention, task engagement). In the next section, we use subjects’ explicit reports during the EL task in order to directly isolate an axis associated with strategy use, and then ask whether the expression of this same axis also predicts better performance in the RL task.

### 1.3 An explicit learning axis extracted during EL predicts performance in the RL task

Several studies suggest that the recruitment of explicit learning processes are associated with better performance during error-based adaptation (de Brouwer et al., 2018), and that such processes reflect task-general executive functions, such as working memory (Anguera et al., 2010). There is evidence also suggesting that such explicit processes can contribute to, and bolster, reinforcement motor learning (Dam et al., 2013; de Brouwer et al., 2021; Codol et al., 2018; Holland et al., 2018). A key prediction from these findings is that the neural recruitment of these task-general, explicit learning processes should be associated with better performance in both EL and RL. Because we do not have a measure of subjects’ explicit strategy use during the RL task, we are unable assay explicit learning in this task directly. However, if explicit learning is supported by task-general processes, then it should be possible to isolate an explicit learning axis using subjects’ explicit reports during the EL task, and show that the expression of this axis during RL predicts performance in that separate task.

To test this, we took the learning epoch of the EL task and computed the partial correlation between subjects’ explicit reports and the mean connectivity of each region with the somatomotor cortex, partialling out subjects’ EL performance scores. These partial correlations thus reflect the effect of explicit strategy use during visuomotor rotation learning, independent of overall performance. For each subject, we then took the inner product between the vector of partial correlations for each region, and the vector containing the mean connectivity of each region with the somatomotor cortex during the learning epoch of the RL task (see Figure 5a). This measure, which we call the *explicit learning score*, reflects the degree to which subjects expressed patterns of functional connectivity associated with explicit learning, as estimated from the EL task, during the RL task.

**Figure 5:**
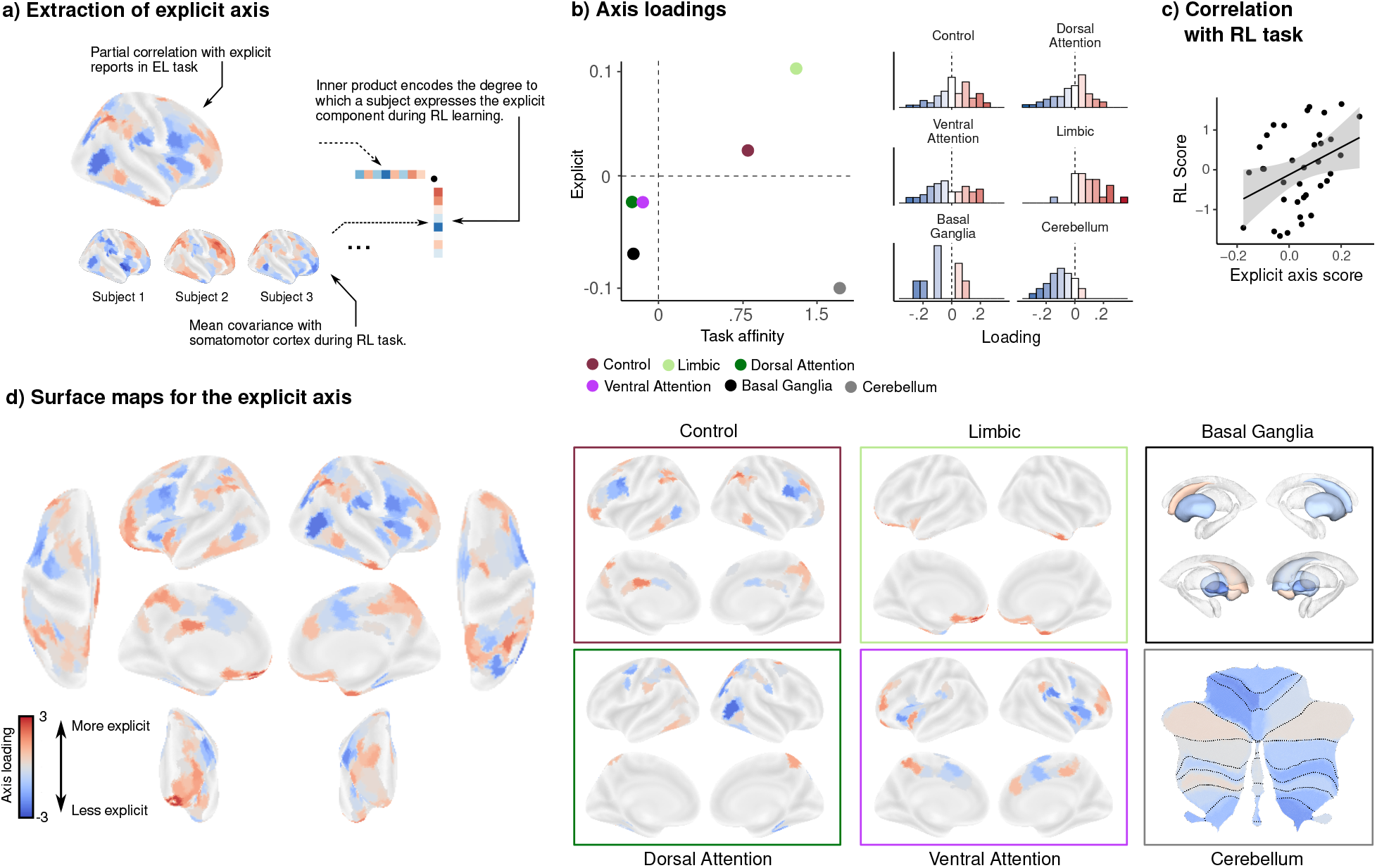
Surface maps for explicit learning axis. **a)** For each ROI, we computed the partial correlation between the mean covariance with the somatomotor cortex during the EL learning epoch, and subjects’ explicit reports. We then projected each subjects’ functional connectivity during the RL learning epoch onto this axis in order to assess the degree to which they expressed this explicit learning axis during the learning phase of the RL task. **b)** (Left) Mean loadings on the explicit and task-affinity axes for each network. (Right) Histograms of explicit axis loadings for individual ROIs in each network. **c)** Greater expression of the explicit learning axis during the RL task was predictive of greater performance. **d)** Surface maps for explicit axis.

Loadings for the explicit axis are shown in Figure 5b. Consistent with evidence that explicit learning acts competitively with implicit adaptation (Albert et al., 2022; Miyamoto et al., 2020), we observed negative correlations between subjects’ explicit reports and functional connectivity in motor regions of the cerebellum. By contrast, we found positive correlations in non-motor cerebellar regions – in particular, the left crus I and II, which have been functionally implicated in more cognitive tasks requiring divided attention, working memory, and mental arithmetic(King et al., 2019). Within the basal ganglia, we also found that correlations were mainly negative, suggesting that explicit learning is associated with reduced basal ganglia connectivity. Within the cortex, loadings in the limbic network were overwhelmingly positive. Loadings in the remaining networks were more mixed, although, within the prefrontal cortex, we observed a broad anterior to posterior gradient in which more anterior regions loaded more positively (see Figure 5d). Notably, when we examined loadings for the explicit axis at the level of subnetworks (see Supplemental Figure 2), we found that correlations were overwhelmingly positive in the Control C network (comprising the posterior cingulate cortex and precuneus), as well as in the control B network, including the dorsolateral prefrontal cortex.

Next, we tested our key prediction – namely, whether the expression of this explicit learning axis, derived from the EL task, would be related to subjects’ performance scores in the RL task (Figure 5c). Consistent with this prediction, we found that the expression of this connectivity axis during the RL task predicted subjects’ RL performance (*r* = .34, *t*_34_ = 2.29, *p* = .028, CI_95_ = [0.04, 0.62]). Together, this suggests that a common explicit component to learning underlies subject-level performance in very different types of motor learning tasks.

## 2 Discussion

Adaptive behaviour can be achieved through both error-based (EL) and reinforcement-based (RL) learning processes, yet the extent to which these two classes of learning recruit distinct versus shared cortical and subcortical neural systems is poorly understood. Here we directly compared, in the same human participants, changes in the connectivity of sensorimotor cortex with distributed areas across cortex, striatum and the cerebellum while individuals learned to adapt their motor commands through either sensory errors (EL) or reward-based feedback (RL). Using supervised dimension reduction techniques, we isolated a dimension of activity, specific to learning, that encoded differences in neural architecture during EL versus RL (the task-affinity axis). We found that while some aspects of the topography of this connectivity axis corresponded with known functional subdivisions in neuroanatomy between EL and RL (e.g., alignment of the cerebellum with EL), there were several other aspects of this topography that differed from our expectations (e.g., alignment of OFC with EL). Notably, we found that the expression of this axis, during EL and RL, was associated with better performance in each task.

Next, to specifically isolate activity patterns associated with explicit learning, which prior work has linked to task-general executive processes (e.g., working memory, attention, and inhibition Anguera et al., 2010; Song and Bédard, 2013; Drummond et al., 2015), we used subjects’ self-reported aiming directions in the EL task to derive a connectivity dimension associated with strategy use (the explicit learning axis). Consistent with our predictions, we found that greater expression of this explicit learning axis in the RL task was associated with greater RL performance. Taken together, our results delineate the cortical and subcortical landscape of activity associated with EL and RL, and demonstrate that sensorimotor learning is supported both by task-specific learning systems, adapted to the unique feedback associated with each task, and task-general cognitive systems, which can be flexibly recruited to bolster performance.

### 2.1 Neural bases of error-based versus reinforcement-based motor learning

As noted above, several topographical features of the task-affinity axis we identified are broadly consistent with proposed functional subdivisions in neuroanatomy supporting EL and RL (Doya, 2000). Specifically, we found that the cerebellum was overwhelmingly EL-aligned, consistent with its established role in implicit learning through sensory prediction errors (Wolpert et al., 2011; Taylor and Ivry, 2014). Our observation that large segments of the dorsal-attention network were EL aligned is likewise consistent with our previous findings that the engagement of the dorsal-attention network is associated with better performance during visuomotor adaptation learning (Areshenkoff et al., 2022). By contrast, we found that the putamen and accumbens were RL-aligned, consistent with their role in reward-based learning (Knowlton et al., 1996). However, we also observed some features of this task-affinity axis that departed from our expectations. For instance, we found that large segments of the superior parietal cortex, particularly in the control and dorsal attention networks, were strongly RL-aligned. Given their well-established role in visually guided action (Andersen et al., 1997), as well as prior studies demonstrating their activity during EL tasks (Anguera et al., 2010; Floyer-Lea and Matthews, 2005; Shadmehr and Holcomb, 1997; Diedrichsen et al., 2005), we had expected these parietal regions to be EL-aligned. Similarly, despite a well-established literature surrounding the role of the limbic system, and orbitofrontal cortex in particular, in RL (Wallis et al., 2007; Rushworth et al., 2007), we found that these areas were instead strongly EL-aligned. Though we can only speculate on the nature of these effects, they may speak to the diversity of processes supported by these higher-order brain areas, and the multiplexed nature of their coding. For example, both of these regions have been strongly implicated, on the one hand, in the encoding of reward associated with different actions (Wallis et al., 2007; Shadlen and Newsome, 2001) and, on the other hand, the spatial encoding of action goals (Yoo et al., 2018; Gottlieb and Snyder, 2010). Thus, it is not obvious a priori how they should align onto our EL and RL tasks. Further, it is recognized that, even in the case of tasks like visuomotor rotation learning, which is often characterized as a classic EL paradigm, performance involves the contribution of multiple implicit learning systems, including those guided by reinforcement (Singh et al., 2019; Taylor and Ivry, 2014; Spampinato and Celnik, 2021; Therrien et al., 2016). Although an ideal comparison of these two classes of learning would involve tasks able to purely isolate EL-versus RL-based processes, it is not obvious how to accomplish this given that, in the context of motor behaviour, these distinct forms of learning are not functionally independent and their neural circuitry cannot be modulated in an on-off fashion (Spampinato and Celnik, 2021). For these reasons, comparisons between these two forms of motor learning, like that done here, often involves the use of tasks that weight differently the contributions of error-based and reinforcement processes required for learning to occur (Spampinato and Celnik, 2021).

Recall that, in order to isolate the task-affinity axis, we needed to disentangle task-specific patterns of activity from activity patterns that were common across both tasks. This involved isolating an additional axis, the task-positive axis, which we found constituted a large increase in cortical and subcortical connectivity from resting-state to task, as well as a relatively smaller increase from baseline to learning. While this pattern of change across these epochs suggests that this connectivity dimension primarily reflects aspects of general task engagement, the fact that its expression reverted to baseline-levels during late learning suggests that it reflects processes particularly at play during early learning. During initial learning, when the sensory and reward prediction errors that drive EL and RL are maximal, it is expected that control, attentional and sensorimotor resources, which all play critical roles in shaping learning (Taylor and Thoroughman, 2007, 2008; Song, 2019), would also be maximally allocated. These resources also likely include the use of task-general strategic processes, but our joint embedding approach did not allow us to disentangle these strategic processes from other features of general task engagement. Instead, our investigation of this issue required a more targeted approach (see next section).

### 2.2 Neural bases of explicit learning

As noted at the outset, error-based visuomotor rotation learning is thought to result from the contribution of different learning components that operate in parallel (Spampinato and Celnik, 2021; Mazzoni and Krakauer, 2006). These include: (1) an implicit learning component that learns by sensory prediction errors, and which is believed to be supported by the cerebellum (Wolpert et al., 2011; Taylor and Ivry, 2014); and, (2) an explicit learning component that is thought to be supported by higher-order association cortex (Taylor and Ivry, 2014) and is associated with better overall learning (de Brouwer et al., 2018, 2021). Although comparably fewer studies have examined the use of reinforcement processes during motor learning (though see Izawa and Shadmehr, 2011; Therrien et al., 2016; Galea et al., 2015; Doyon et al., 2009), there is emerging behavioural evidence that a similar parallel system architecture may also govern RL, and that EL and RL are both supported by a common explicit learning component (Holland et al., 2018; de Brouwer et al., 2021).

To study the neural basis of this explicit learning component directly, we isolated a dimension of activity (the explicit learning axis) that covaried with subjects’ explicit reports in the EL task, as these provided a direct assay of explicit learning (Taylor et al., 2014). Our analysis revealed that explicit knowledge of the rotation in the EL task was associated with greater functional connectivity between somatomotor cortices and, most consistently, regions in the limbic and control B/C networks. The latter control network comprises the posterior cingulate cortex, which has been implicated in the top-down selection of response strategies spanning multiple trials (Pearson et al., 2009, 2011); and the precuneus, which has been implicated in the voluntary deployment of spatial attention (Ogiso et al., 2000; Wenderoth et al., 2005; Krumbholz et al., 2009). Control network B prominently includes the dorsolateral prefrontal cortex, which in particular has been implicated in the use of spatial information for action selection (see Hoshi, 2006, for a review). This explicit learning axis thus incorporates a variety of regions commonly implicated in both general strategic processing, as well as the strategic use of spatial information specifically. It is thus intuitive that the interaction of these particular regions with somatomotor cortex should be associated with the use of use of explicit strategies during EL.

We reasoned that if this explicit learning axis does indeed reflect, at least in part, task-general strategic processes, then the expression of this axis during the separate RL task should predict better RL performance. Our findings not only confirmed this prediction, but they also bolstered our interpretation of this axis as reflecting the explicit learning component; namely, the fact that this axis constituted a concomitant increase in anterior prefrontal connectivity and decrease in anterior (motor) cerebellar connectivity is consistent with behavioural work showing that an increase in explicit learning decreases the contribution of implicit learning, by depriving it of the errors required to drive gradual adaptation (Albert et al., 2022; Miyamoto et al., 2020; McDougle et al., 2015; Kim et al., 2019; Leow et al., 2020; Avraham et al., 2021).

### 2.3 Methodological Considerations

Our approach, using supervised PCA, allowed us to derive dimensions of activity that have precise relationships to the structure of our task, including connectivity axes distinguishing between specific groups of scans and/or epochs of the task, and correlating with specific patterns of behaviour. As a large body of research suggests that the dominant patterns of variability in functional connectivity are related either to the gross topographical properties of functional brain organization (Margulies et al., 2016), or to static subject-level differences unrelated to task performance (Gratton et al., 2018), our unique approach – combined with our covariance centering – allowed us to speak more precisely about the activity patterns related to learning in each task.

For practical reasons, we restricted our analysis to bipartite networks describing functional interactions between the somatomotor cortex and a variety of regions implicated in implicit (cerebellum and striatum) and explicit (cortical) motor learning processes. We believe this is a sensible restriction, as the motor cortex serves as the primary output pathway in the motor system, and it is functional interactions with this system, in particular, that are most likely to result in modifications to behaviour (i.e., learning). However, a limitation of this targeted approach is that we cannot speak to connectivity changes occurring within and between the remaining regions that may also relate to behaviour. We felt this was a reasonable trade-off, as it is often difficult to interpret effects across whole-brain networks without either resorting to coarse summary statistics, or requiring a large number of individual comparisons.

Our derivation of the explicit learning axis was based on subjects’ self-reported aim direction during the report block. Although this measure is well validated, and correlates well with features of performance typically associated with explicit learning (Taylor et al., 2014; de Brouwer et al., 2018), we do note that some subjects expressed confusion about the nature of the report trials. As a result, some subjects whom we have characterized as fully implicit in actuality may have simply failed to understand the task required of them (thus leaving the report dial pointing at the target location). We believe this explains the large performance variability in subjects reporting zero explicit knowledge (Figure 2c). Nevertheless, given both the existing literature (Taylor et al., 2014; de Brouwer et al., 2018, 2021) and the fact that we observe a clear relationship between performance and reported aim direction among subjects reporting a non-zero re-aiming direction, we believe that our reports provide a valid, albeit noisy, metric of subjects’ explicit strategy use.

Finally, whereas previous descriptions of the cortico-subcortical pathways governing EL and RL have often been reliant on indirect comparisons assembled across different studies, species and motor-related tasks, our approach; studying both forms of learning within the same subjects, and jointly embedding both tasks into a common neural space, allowed us to speak directly to the differences and commonalities in low-dimensional network structure associated with each task. We believe it is this repeated-measures approach that has afforded us a unique view into cortical landscape governing the two major forms of motor learning.

## 3 Methods

### 3.1 Participants

Fourty-six right-handed subjects (27 females, aged 18-28 years, mean age: 20.3 years) participated in three separate testing sessions, each spaced approximately 1 week apart: the first, an MRI-training session, was followed by two subsequent MRI experimental sessions. Of these 46 subjects, one subject was excluded due of excessive head motion in the MRI scanner (motion greater than 2 mm or 2° rotation within a single scan). One was encluded due interuption of the scan during the learning phase of the RL task. Five subjects were excluded due to poor behavioural performance in one or both tasks (4 of these participants were excluded because >25% of trials were not completed within the maximum trial duration; and one because >20% of trials had missing data due to insufficient pressure of the fingertip on the MRI-compatible tablet). Finally, three were excluded due to a failure to properly perform the RL task; these subjects did not trace the visible rightward path during the baseline phase, but rather continued to trace a straight line for the entire duration of the task, suggesting that they did not attend to the task, and making it difficult to interpret their scores during the learning block. We used the remaining 36 participants for analysis. Right-handedness was assessed with the Edinburgh handedness questionnaire (Oldfield, 1971). Participant informed was obtained before beginning the experimental protocol. The Queen’s University Research Ethics Board approved the study and it was conducted in coherence to the principles outlines in the Canadian Tri-Council Policy Statement on Ethical Conduct for Research Involving Humans and the principles of the Deceleration of Helsinki (1964).

### 3.2 Procedure

In the first session, participants took part in an MRI-training session inside a mock (0 T) scanner, that was made to look and sound like a real MRI scanner. We undertook this training session to (1) introduce participants to the two separate experimental tasks to be subsequently performed in the MRI scanner (a Reinforcement Learning (RL) task and an Error-based Learning (EL) task; see below for details on the two separate tasks), (2) ensure that subjects obtained baseline performance levels on those two tasks, and (3) ensure that participants could remain still without feeling claustrophobic for 1 hr. To equate baseline performance levels across participants, subjects performed approximately 80 trials per task. To help ensure minimal participant movement in the mock scanner, we monitored subjects’ head movement by attaching, via tape, a Polhemus sensor to each subject’s forehead (Polhemus, Colchester, Vermont). This allowed a real-time read-out of subject head displacement in the three axes of translation and rotation (6 dimensions total). Whenever subjects’ head translation and/or rotation reached 0.5 mm or 0.5° rotation (within some velocity criterion), subjects received an unpleasant auditory tone, delivered through a speaker system located near the head. All the subjects learned to constrain their head movement via this auditory feedback. Following this first training session, subjects then subsequently participated in RL and EL tasks, respectively (see below for details).

### 3.3 Apparatus

During testing in the mock (0 T) scanner, subjects performed hand movements that were directed towards a target by applying finger tip pressure on a digitizing touchscreen tablet (Wacom Intuos Pro M tablet). During the actual MRI testing sessions, subjects used an MRI-compatible digitizing tablet (Hybridmojo LLC, CA, USA). In both the mock and real MRI scanner the target and cursor stimuli were rear-projected with an LCD projector (NEC LT265 DLP projector, 1024 x 768 resolution, 60 Hz refresh rate) onto a screen mounted behind the participant. The stimuli on the screen were viewed through a mirror fixated on the MRI coil directly above the participants’ eyes, thus preventing the participant from being able to see their hand.

### 3.4 Reinforcement Learning (RL) task

In this task, participants were trained, through reward-based trial-and-error learning, to produce finger movement trajectories with a specific (unseen) shape. Specifically, subjects were instructed to repeatedly trace, without visual cursor feedback of their actual finger paths, a subtly curved path displayed on the screen (the visible path, Figure 1b, black trace). Participants were told that, following each trial, they would receive a score based on how accurately they traced the visible path (and instructed to maximize points across trials). However, unbeknownst to them, they actually received points based on how well they traced the mirror-image path (the target path, Figure 1b, dashed black trace). Critically, because participants received no visual feedback about their actual finger trajectories or the ‘rewarded’ shape, they could not use error-based information to guide learning. Our task was a slight modification on the task developed by (Wu et al., 2014).

Each trial started with the participant moving the cursor (3 mm radius cyan circle) into the start position (4 mm radius white circle) at the bottom of the screen by sliding the index finger on the tablet. The cursor was only visible when it was within 30 mm of the start position. After the cursor was held within the start position for 0.5 s, the cursor disappeared and a curved path and a target appeared on the screen. The target was a horizontal red line (30 x 1 mm) that appeared 60 mm above the start position. The path connected the start and target position and had the shape of a half sine wave with an amplitude of 0.15 times the target distance. Participants were instructed to trace the curved path. When the cursor reached the target distance, the target changed colour from red to green to indicate that the trial was completed. Participants did not receive feedback about the position of their cursor.

In the baseline block, participants did not receive feedback about their performance. In the learning block, participants were rewarded 0 to 100 points after reaching the target, and participants were instructed to do their best to maximize this score (the points were displayed as text centrally on the screen). They were told that to increase the score, they had to trace the line more accurately. Each trial was terminated after 4.5 s, independent of whether the cursor had reached the target. After a delay of 1.5 s, allowing time to save the data, the next trial started with the presentation of the start position.

To calculate the score on each trial in the learning block, the x position of the cursor was interpolated at each cm displacement from the start position in the y direction (i.e., at exactly 10, 20, 30, 40, 50 and 60 mm). For each of the six y positions, the absolute distance between the interpolated x position of the cursor and the x position of the rewarded path was calculated. The sum of these errors was scaled by dividing it by the sum of errors obtained for a half cycle sine-shaped path with an amplitude of 0.5 times the target distance, and then multiplied by 100 to obtain a score ranging between 0 and 100. The scaling worked out so that a perfectly traced line would result in an imperfect score of 40 points. This scaling was chosen on the basis of extensive pilot testing in order to achieve a variety of subject performance, and to ensure that subjects still received informative score feedback when tracing in the vicinity of the visible trajectory.

During the participant training session in the mock MRI scanner (i.e., 1-week prior to the MRI testing session), participants only performed a practice block in which they traced a straight line with (40 trials) and then without (40 trials) visual feedback of the position of the cursor during the reach. This training session exposed participants to several key features of the task (e.g., use of the touchscreen tablet, trial timing, presence and removal of cursor feedback), allowed us to establish adequate performance levels, but importantly did not allow for any reward-based learning to take place.

At the beginning of the MRI testing session, but prior to the first scan being collected, participants re-acquainted themselves with the RL task by first performing a practice block in which they traced a straight line with (40 trials) and then without (40 trials) visual feedback of the position of the cursor during the reach. Next, we collected an anatomical scan, following by a DTI scan, followed by a resting-state fMRI scan. During the latter resting-state scans, participants were instructed to rest with their eyes open while fixating on a central cross location presented on the screen. Following this, participants performed the RL task, which consisted of two separate experimental runs without visual feedback of the cursor: (1) a baseline block of 70 trials in which they attempted to trace the curved path and no score was provided, and (2) a separate learning block of 200 trials in which participants were instructed to maximize their score shown at the end of each trial.

### 3.5 Error-based Learning (EL) task

To probe EL, we used the well-characterized visuomotor rotation (VMR) paradigm (Krakauer, 2009). During the VMR task, participants performed blocks of trials in which they first used their left, and subsequently their right, index finger to perform center-out target-directed movements. After these baseline trials, we applied a 45° clockwise (CW) rotation to the viewed cursor, allowing investigation of learning with the right hand. Following this, we briefly assessed participants’ re-aiming strategy associated with their learning. Finally, to permit investigation of intermanual transfer, subjects switched hands and performed the same VMR task with their untrained left hand.

Each trial started with the participant moving the cursor (3 mm radius cyan circle) into the start position (4 mm radius white circle) in the centre of the screen by sliding the index finger on the tablet. To guide the cursor to the start position, a ring centred around the start position indicated the distance between the cursor and the start position. The cursor became visible when its centre was within 8 mm of the centre of the start position. After the cursor was held within the start position for 0.5 s, a target (5 mm radius red circle) was shown on top of a grey ring with a radius of 60 mm (i.e., the target distance) centred around the start position. The target was presented at one of eight locations, separated by 45° (0, 45, 90, 135, 180, 225, 270 and 315°), in randomized bins of eight trials. Participants were instructed to hit the target with the cursor by making a fast finger movement on the tablet. They were instructed to ‘slice’ the cursor through the target to minimize online corrections during the reach. If the movement was initiated (i.e., the cursor had moved fully out of the start circle) before the target appeared, the trial was aborted and a feedback text message “Too early” appeared centrally on the screen. In trials with correct timing, the cursor was visible during the movement to the ring and then became stationary for one second when it reached the ring, providing the participant with visual feedback of their endpoint reach error. If any part of the stationary cursor overlapped with any part of the target, the target coloured green to indicate a hit. Each trial was terminated after 4.5 s, independent of whether the cursor had reached the target. After a delay of 1.5 s, allowing time to save the data, the next trial started with the presentation of the start position.

During the participant training session in the mock MRI scanner (i.e., 2-weeks prior to the VMR MRI testing session), participants performed a practice block of 80 trials (40 trials with with their left hand and 40 trials with their right hand) with veridical feedback (i.e., no rotation was applied to the cursor). As with the RL task, this training session exposed participants to several key features of the task (e.g., use of the touchscreen tablet, trial timing, used of cursor feedback to correct for errors) and allowed us to establish adequate performance levels.

At the beginning of the MRI testing session, but prior to the first scan being collected, participants re-acquainted themselves with the EL task by performing 80 practice trials with veridical cursor feedback (40 trials with each hand). Next, we collected an anatomical scan, followed by a consecutive series of 4 fMRI experimental runs. The first run included baseline blocks of 64 trials with veridical cursor feedback that were performed with the left hand. The second run was identical, except was performed with the right hand. The third run included a rotation block of 160 trials in which feedback of the cursor during the reach was rotated clockwise by 45°, and was performed with the right hand. Importantly, following this and prior to beginning the last (fourth) fMRI experimental run, participants were asked to report their strategic aiming direction (over 16 trials). In these trials, a line between the start and target position appeared on the screen at the start of each trial. Participants were asked to use a separate MRI joystick (Current Designs, Inc.) positioned at their left hip to rotate the line to the direction that they would aim their finger movement in for the cursor to hit the target, and click the button on the joystick box when satisfied. Following the button click, the trial proceeded as a normal reach trial. These 16 “report” trials were followed by 16 normal rotation trials. As several subjects expressed confusion about the nature of the report trials, often failing to adjust the line at all during the first few trials, we discarded the first 8 report trials for all subjects, and calculated the mean aim direction using the final 8 trials. Note that we had subjects perform these report trials following learning (as in de Brouwer et al., 2021) given our prior behavioural work showing that the inter-mixing of report trials during learning can lead to more participants adopting an explicit, re-aiming strategy (de Brouwer et al., 2018, 2021), thereby distorting participants’ learning curves. Following this, we resumed fMRI testing and the fourth run included a rotation block of 80 trials that was identical, except was performed with the left hand (allowing investigation of intermanual transfer; this data was not analyzed here). Participants were not informed about nature or presence of the visuomotor rotation before or during the experiment.

### 3.6 Behavioral data analysis

#### 3.6.1 EL task preprocessing

Trials in which the reach was initiated before the target appeared (4% of trials) or in which the cursor did not reach the target within the time limit (5% of trials) were excluded from the offline analysis of hand movements. As insufficient pressure on the touchpad resulted in a default state in which the cursor was reported as lying in the top left corner of the screen, we excluded trials in which the cursor jumped to this position before reaching the target region (2% of trials). We then applied a conservative threshold on the movement and reaction times, removing the top .05% of trials across all subjects. As the EL required the subject to determine the target location prior to responded, we also set a lower threshold of 100 ms on the reaction time.

We then calculated the angle difference between the target position and the last sample before the cursor exceeded the target circle. This was then converted to an adaptation score by subtracting the angle of the rotation on each trial (0° during baseline, and 45° during learning) in order to derive the actual aim direction on each trial. For consistency with the RL task, we flipped the sign of this score so that positive values always reflected greater performance.

#### 3.6.2 RL task preprocessing

Trials in which the cursor did not reach the target within the time limit were excluded from the offline analysis of hand movements (1% of trials). As insufficient pressure on the touchpad resulted in a default state in which the cursor was reported as lying in the top left corner of the screen, we excluded trials in which the cursor jumped to this position before reaching the target region (2% of trials). We then applied a conservative threshold on the movement and reaction times, removing the top .05% of trials across all subjects. As the RL task did not involve response discrimination, we did not set a lower threshold on these variables.

#### 3.6.3 Functional PCA of subject behavioral data

All subject behavioral data were averaged over 8 trial blocks. In the EL task, this corresponded to one presentation of each target location. For consistency between tasks, an identical window size was chosen for the RL task. We represented individual learning curves as functional data using a cubic spline basis with smoothing penalty estimated by generalized cross-validation (Härdle, 1990), and for each task separately performed functional PCA (CITE) in order to extract components capturing the dominant patterns of variability in subject performance. In both tasks, the top component described overall learning, and explained a majority of the variability (70%) in performance. Spline smoothing and fPCA were performed using the R package fda (Ramsay et al., 2022), and example code is provided in a publically available repository (see Data and Code Availability statement).

#### 3.6.4 Bayesian multivariate regression model

Our model of subjects’ joint performance in the EL and RL tasks (see Figure 1) is essentially a bivariate linear regression model with correlated errors, allowing subjects’ reported aim direction in the EL task to affect both the mean performance in each task, as well as the error correlation. The effect of the explicit report is assumed to be linear on the scale of the Fisher transformed correlation.

Let **y**_EL_ and **y**_RL_ be vectors of subjects’ EL and RL component scores, respectively, and let **x**_Rep_ be subjects mean reported aim direction during the EL report block. We fit a multivariate regression model

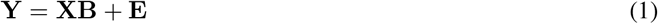

where **Y** = [**y**_EL_ **y**_RL_] and **X** = [**1 x**_Rep_] is the design matrix. We assume that the errors *ϵ_i_* = [*ϵ*_EL_*i*__ *ϵ*_RL_*i*__] are independent and jointly normal, with covariance

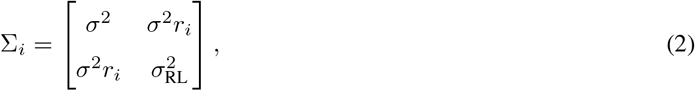

where *r_i_* is the correlation between **e**_EL_*i*__. and **e**_RL_*i*__. The Fisher transform of *r_i_* is allowed to vary as a linear function of **x**_Rep_, so that

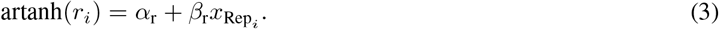

We place weakly informative standard normal priors over the coefficients, and half-Cauchy priors over the covariance parameters 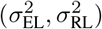. The full model is thus

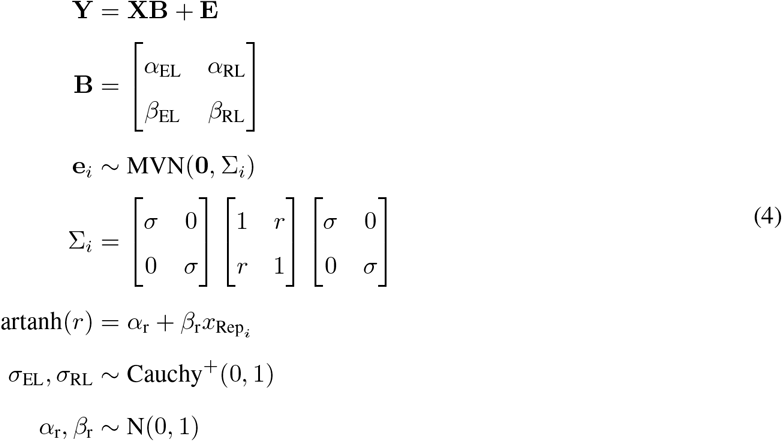

The mode was fit by Hamiltonian Monte-Carlo in RStan (Stan Development Team, 2022) using 4 chains of 4000 samples, of which the first 1000 were discarded as burn-in. Chains were inspected visually, and for all parameters we observed 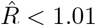.

To visualize the model predictions, we used the posterior mean parameters estimates. For a given explicit report *x*_Rep_*i*__, the predicted distribution of EL and RL scores is then

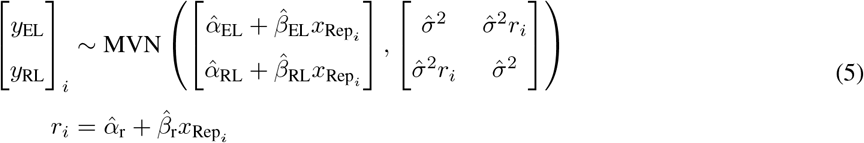

Where hats denote posterior mean estimates. The predictions are illustrated in Figure 2d (bottom right) as confidence ellipses representing a Mahalanobis distance of 1 from the mean. Code and example data for implementing the model are provided in a Github repository (see data and code availability statement).

### 3.7 MRI Acquisition

Participants were scanned using a 3-Tesla Siemens TIM MAGNETOM Trio MRI scanner located at the Centre for Neuroscience Studies, Queen’s University (Kingston, Ontario, Canada). Functional MRI volumes were acquired using a 32-channel head coil and a T2*-weighted single-shot gradient-echo echo-planar imaging (EPI) acquisition sequence (time to repetition (TR) = 2000 ms, slice thickness = 4 mm, in-plane resolution = 3 mm x 3 mm, time to echo (TE) = 30 ms, field of view = 240 mm x 240 mm, matrix size = 80 x 80, flip angle = 90°, and acceleration factor (integrated parallel acquisition technologies, iPAT) = 2 with generalized auto-calibrating partially parallel acquisitions (GRAPPA) reconstruction. Each volume comprised 34 contiguous (no gap) oblique slices acquired at a 30° caudal tilt with respect to the plane of the anterior and posterior commissure (AC-PC), providing whole-brain coverage of the cerebrum and cerebellum. Each of the task-related scans included an additional 8 imaging volumes at both the beginning and end of the scan. On average, each of the MRI testing sessions lasted 2 hrs.

At the beginning of the RL MRI testing session, a T1-weighted ADNI MPRAGE anatomical was also collected (TR = 1760 ms, TE = 2.98 ms, field of view = 192 mm x 240 mm x 256 mm, matrix size = 192 x 240 x 256, flip angle = 9°, 1 mm isotropic voxels). Next, we collected a resting-state scan, wherein 300 imaging volumes were acquired. For the baseline and learning scans during RL testing, 222 and 612 imaging volumes were acquired, respectively.

At the beginning of the EL MRI testing session, we gathered high-resolution whole-brain T1-weighted (T1w) and T2-weighted (T2w) anatomical images (in-plane resolution 0.7 x 0.7 mm2; 320 x 320 matrix; slice thickness: 0.7 mm; 256 AC-PC transverse slices; anterior-to-posterior encoding; 2 x acceleration factor; T1w TR 2400 ms; TE 2.13 ms; flip angle 8°; echo spacing 6.5 ms; T2w TR 3200 ms; TE 567 ms; variable flip angle; echo spacing 3.74 ms). These protocols were selected on the basis of protocol optimizations designed by Filippini et al. (2014). Following this, for the baseline, learning and transfer functional scans, 204, 492 and 252 imaging volumes were acquired, respectively.

### 3.8 fMRI Preprocessing

Results included in this manuscript come from preprocessing performed using *fMRIPrep* 1.4.1 (Esteban et al. (2018b); Esteban et al. (2018a); RRID:SCR_016216), which is based on *Nipype* 1.2.0 (Gorgolewski et al. (2011); Gorgolewski et al. (2018); RRID:SCR_002502).

**Anatomical data preprocessing** A total of 2 T1-weighted (T1w) images were found within the input BIDS dataset. All of them were corrected for intensity non-uniformity (INU) with `N4BiasFieldCorrection` (Tustison et al., 2010), distributed with ANTs 2.2.0 (Avants et al., 2008, RRID:SCR_004757). The T1w-reference was then skull-stripped with a *Nipype* implementation of the `antsBrainExtraction.sh` workflow (from ANTs), using OASIS30ANTs as target template. Brain tissue segmentation of cerebrospinal fluid (CSF), white-matter (WM) and gray-matter (GM) was performed on the brain-extracted T1w using `fast` (FSL 5.0.9, RRID:SCR_002823, Zhang et al., 2001). A T1w-reference map was computed after registration of 2 T1w images (after INU-correction) using `mri_robust_template` (FreeSurfer 6.0.1, Reuter et al., 2010). Brain surfaces were reconstructed using `recon-all` (FreeSurfer 6.0.1, RRID:SCR_001847, Dale et al., 1999), and the brain mask estimated previously was refined with a custom variation of the method to reconcile ANTs-derived and FreeSurfer-derived segmentations of the cortical gray-matter of Mindboggle (RRID:SCR_002438, Klein et al., 2017). Volume-based spatial normalization to two standard spaces (MNI152NLin6Asym, MNI152NLin2009cAsym) was performed through nonlinear registration with `antsRegistration` (ANTs 2.2.0), using brain-extracted versions of both T1w reference and the T1w template. The following templates were selected for spatial normalization: *FSL’s MNI ICBM 152 non-linear 6th Generation Asymmetric Average Brain Stereotaxic Registration Model* [Evans et al. (2012), RRID:SCR_002823; TemplateFlow ID: MNI152NLin6Asym], *ICBM 152 Nonlinear Asymmetrical template version 2009c* [Fonov et al. (2009), RRID:SCR_008796; TemplateFlow ID: MNI152NLin2009cAsym].
**Functional data preprocessing** For each of the 5 BOLD runs found per subject (across all tasks and sessions), the following preprocessing was performed. First, a reference volume and its skull-stripped version were generated using a custom methodology of *fMRIPrep*. The BOLD reference was then co-registered to the T1w reference using `bbregister` (FreeSurfer) which implements boundary-based registration (Greve and Fischl, 2009). Co-registration was configured with nine degrees of freedom to account for distortions remaining in the BOLD reference. Head-motion parameters with respect to the BOLD reference (transformation matrices, and six corresponding rotation and translation parameters) are estimated before any spatiotemporal filtering using `mcflirt` (FSL 5.0.9, Jenkinson et al., 2002). BOLD runs were slice-time corrected using `3dTshift` from AFNI 20160207 (Cox and Hyde, 1997, RRID:SCR_005927). The BOLD time-series, were resampled to surfaces on the following spaces: *fsaverage*. The BOLD time-series (including slice-timing correction when applied) were resampled onto their original, native space by applying a single, composite transform to correct for head-motion and susceptibility distortions. These resampled BOLD time-series will be referred to as *preprocessed BOLD in original space*, or just *preprocessed BOLD*. The BOLD time-series were resampled into several standard spaces, correspondingly generating the following *spatially-normalized, preprocessed BOLD runs:* MNI152NLin6Asym, MNI152NLin2009cAsym. First, a reference volume and its skull-stripped version were generated using a custom methodology of *fMRIPrep*. Automatic removal of motion artifacts using independent component analysis (ICA-AROMA, Pruim et al., 2015) was performed on the *preprocessed BOLD on MNI space* time-series after removal of non-steady state volumes and spatial smoothing with an isotropic, Gaussian kernel of 6mm FWHM (full-width half-maximum). Corresponding “non-aggresively” denoised runs were produced after such smoothing. Additionally, the “aggressive” noise-regressors were collected and placed in the corresponding confounds file. Several confounding time-series were calculated based on the *preprocessed BOLD:* framewise displacement (FD), DVARS and three region-wise global signals. FD and DVARS are calculated for each functional run, both using their implementations in *Nipype* (following the definitions by Power et al., 2014). The three global signals are extracted within the CSF, the WM, and the whole-brain masks. Additionally, a set of physiological regressors were extracted to allow for component-based noise correction (*CompCor*, Behzadi et al., 2007). Principal components are estimated after high-pass filtering the *preprocessed BOLD* time-series (using a discrete cosine filter with 128s cut-off) for the two *CompCor* variants: temporal (tCompCor) and anatomical (aCompCor). tCompCor components are then calculated from the top 5% variable voxels within a mask covering the subcortical regions. This subcortical mask is obtained by heavily eroding the brain mask, which ensures it does not include cortical GM regions. For aCompCor, components are calculated within the intersection of the aforementioned mask and the union of CSF and WM masks calculated in T1w space, after their projection to the native space of each functional run (using the inverse BOLD-to-T1w transformation). Components are also calculated separately within the WM and CSF masks. For each CompCor decomposition, the *k* components with the largest singular values are retained, such that the retained components’ time series are sufficient to explain 50 percent of variance across the nuisance mask (CSF, WM, combined, or temporal). The remaining components are dropped from consideration. The head-motion estimates calculated in the correction step were also placed within the corresponding confounds file. The confound time series derived from head motion estimates and global signals were expanded with the inclusion of temporal derivatives and quadratic terms for each (Satterthwaite et al., 2013). Frames that exceeded a threshold of 0.5 mm FD or 1.5 standardised DVARS were annotated as motion outliers. All resamplings can be performed with *a single interpolation step* by composing all the pertinent transformations (i.e. head-motion transform matrices, susceptibility distortion correction when available, and co-registrations to anatomical and output spaces). Gridded (volumetric) resamplings were performed using `antsApplyTransforms` (ANTs), configured with Lanczos interpolation to minimize the smoothing effects of other kernels (Lanczos, 1964). Non-gridded (surface) resamplings were performed using `mri_vol2surf` (FreeSurfer).

Many internal operations of *fMRIPrep* use *Nilearn* 0.5.2 (Abraham et al., 2014, RRID:SCR_001362), mostly within the functional processing workflow. For more details of the pipeline, see the section corresponding to workflows in *fMRIPrep’s* documentation.

#### 3.8.1 ROI extraction

Regions of interest were identified using separate parcellations for the cerebral cortex, striatum, and cerebellum. We extracted all cerebellar regions from the atlas of Diedrichsen (Diedrichsen, 2006). Striatal regions; including the caudate, putamen, accumbens, and pallidum, were extracted from the Harvard Oxford atlas (Frazier et al., 2005; Makris et al., 2006). Cortical regions were extracted from the 400 region Schaefer parcellation (Schaefer et al., 2018). Specifically, we extracted left-hemispheric ROIs in the Somatomotor A network (corresponding to the hand area in the somatomotor cortices), as well as bilateral ROIs belonging to the control, limbic, dorsal attention, and ventral attention networks. We then extracted the standardized mean BOLD signal within each ROI.

### 3.9 Neuroimaging data analysis

#### 3.9.1 Covariance estimation and centering

For each subject, we used the Ledoit and Wolf shrinkage estimator (Ledoit and Wolf, 2004) to derive covariance matrices for the resting state scan, as well as periods of equivalent length (100 imaging volumes) during baseline, and the beginning and end of the learning blocks of each task. For the the resting state and baseline blocks, these 100 volumes were extracted at equal intervals throughout the scans.

To center the covariance matrices, we took the approach advocated by Zhao et al. (2018), which leverages the natural geometry of the space of covariance matrices. We have implemented many of the computations required to replicate the analysis in an publically available R package **spdm**, which is freely available from a Git repository at https://github.com/areshenk-rpackages/spdm.

The procedure is as follows. Letting *S_ij_* denote the *j*’th covariance matrix for subject *i*, we computed the grand mean covariance 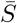 over all subjects using the fixed-point algorithm described by Congedo et al. (2017), as well as subject means 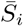. We then projected the each covariance matrix *S_ij_* onto the tangent space at 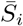 to obtain a tangent vector

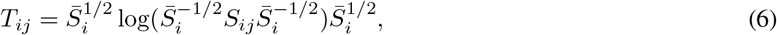

where log denotes the matrix logarithm. The tangent vector *T_ij_* then encodes the *difference* in covariance between the covariance *S_ij_* and the subject mean 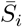. We then transported each tangent vector to the grand mean 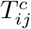 using the transport proposed by Zhao et al. (2018), obtaining a centered tangent vector 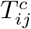 given by

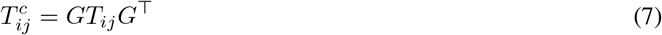

where 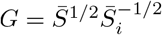. This centered tangent vector now encodes the same difference in covariance, but now expressed relative to the mean resting state scan. Finally, we projected each centered tangent vector back onto the space of covariance matrices, to obtain the centered covariance matrix

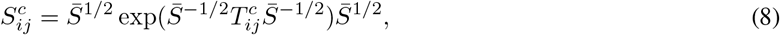

where exp denotes the matrix exponential.

In order to visualize the effects of centering, used uniform manifold approximation (UMAP; McInnes et al., 2018) on the matrix of pairwise geodesic distances between covariance matrices. This embedding revealed that centering was successful in eliminating static differences in covariance, and in isolating the effect of task condition.

#### 3.9.2 Supervised PCA

After centering, we projected the centered covariance matrices onto the tangent space around the grand mean covariance. We then extracted slices of these tangent vectors corresponding to the connectivity between the regions in the Somatomotor A cortical network, and the remaining cortical and subcortical ROIs. For each of the latter ROIs, we computed the average connectivity across all regions in the Somatomotor network, obtaining, for each subject and each epoch, a vector encoding the mean connectivity of each region with the somatomotor cortex.

In order to extract components corresponding to task-general and task-specific functional connectivity, we performed supervised PCA on these vectors, using the dual formulation of Barshan et al. (2011) which allows the specification of a target kernel matrix for which the estimated principal component attempt to minimize prediction error. The target variables for our sPCA comprised two respective feature vectors **y**_*g*_ and **y**_*s*_:

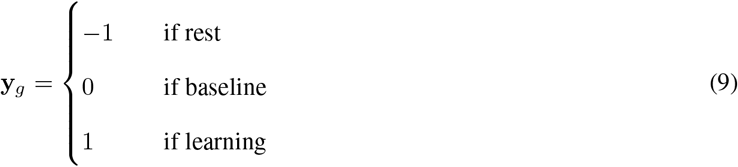

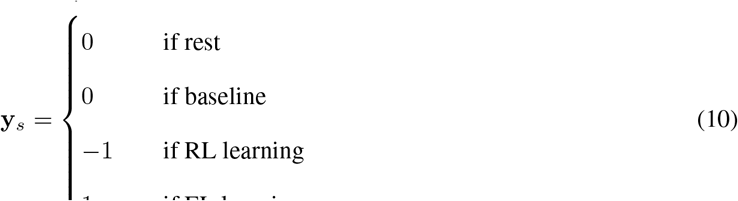

The target of the sPCA was then the kernel matrix

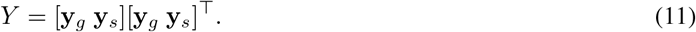

Code implementing the procedure is available in the linked repository (see Data and code availability statement).

## Acknowledegments

C.N.A. was supported by a Natural Sciences and Engineering Research Council (NSERC) graduate award. J.P.G. was also supported by a NSERC Discovery Grant (RGPIN-2017-04684), Canadian Institutes of Health Research Grant (MOP126158), and Botterell Foundation Award, as well as funding from the Canadian Foundation for Innovation (35559). The authors would like to thank Martin York and Sean Hickman for technical assistance, and Don O’Brien for assistance with data collection.

## Data availability

Behavioral and imaging data are archived at Queens University and are available from the authors upon reasonable request.

## Code availability

Imaging data were preprocessed using fmriPrep 1.4.0, which is open source and freely available. Operations on covariance matrices, including estimation and centering, were performed using the R package spdm, which is freely available in a repository at https://github.com/areshenk-rpackages/spdm. Tutorial code and data for implementing the centering procedure, supervised PCA, and functional PCA are hosted in a GitHub repository at https://github.com/areshenk-opendata/2022-elrl-subspace.

---

**Supplemental Figure 1:**
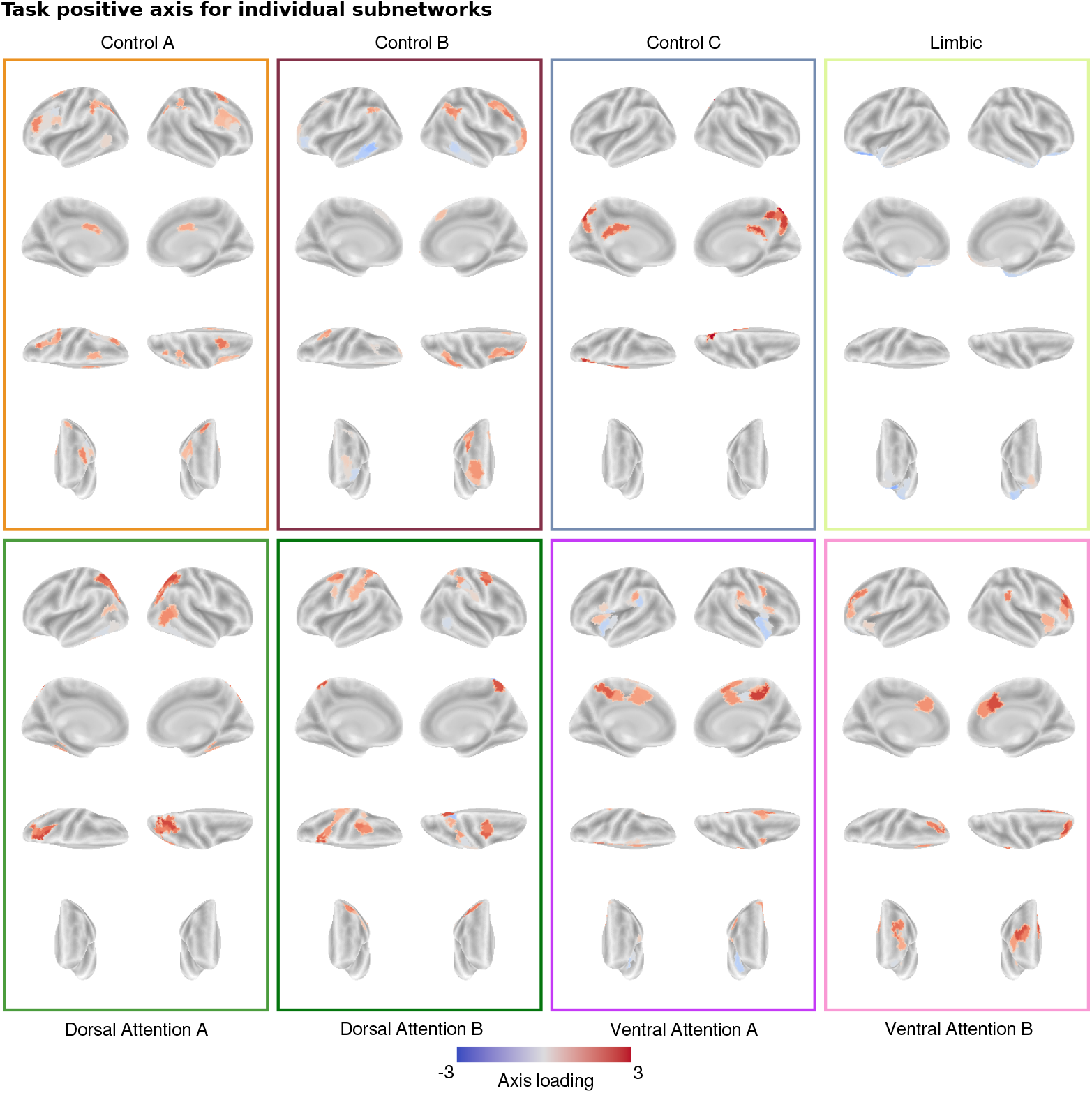
Task positive axis for individual subnetworks.

**Supplemental Figure 2:**
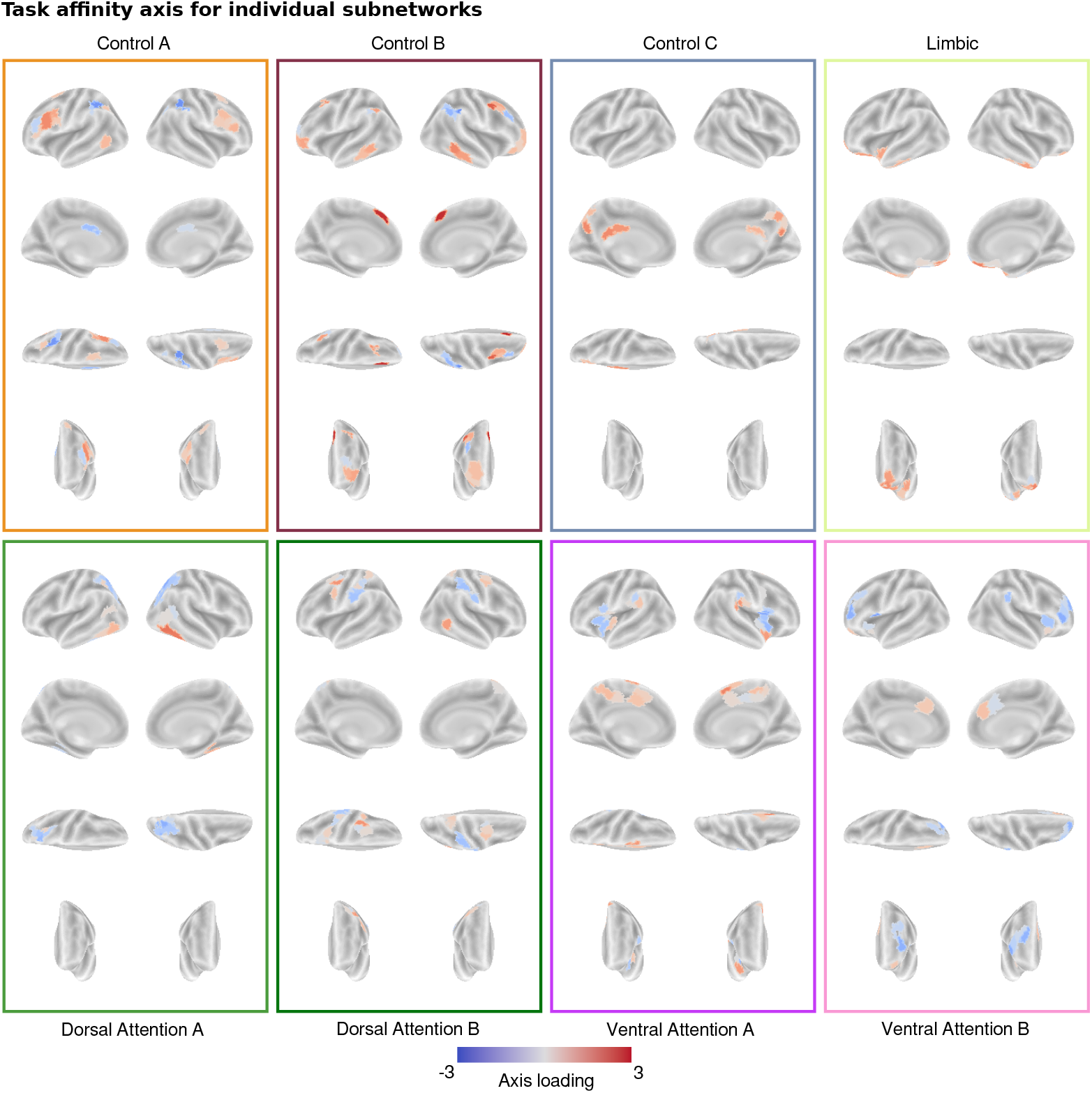
Task affinity axis for individual subnetworks.

**Supplemental Figure 3:**
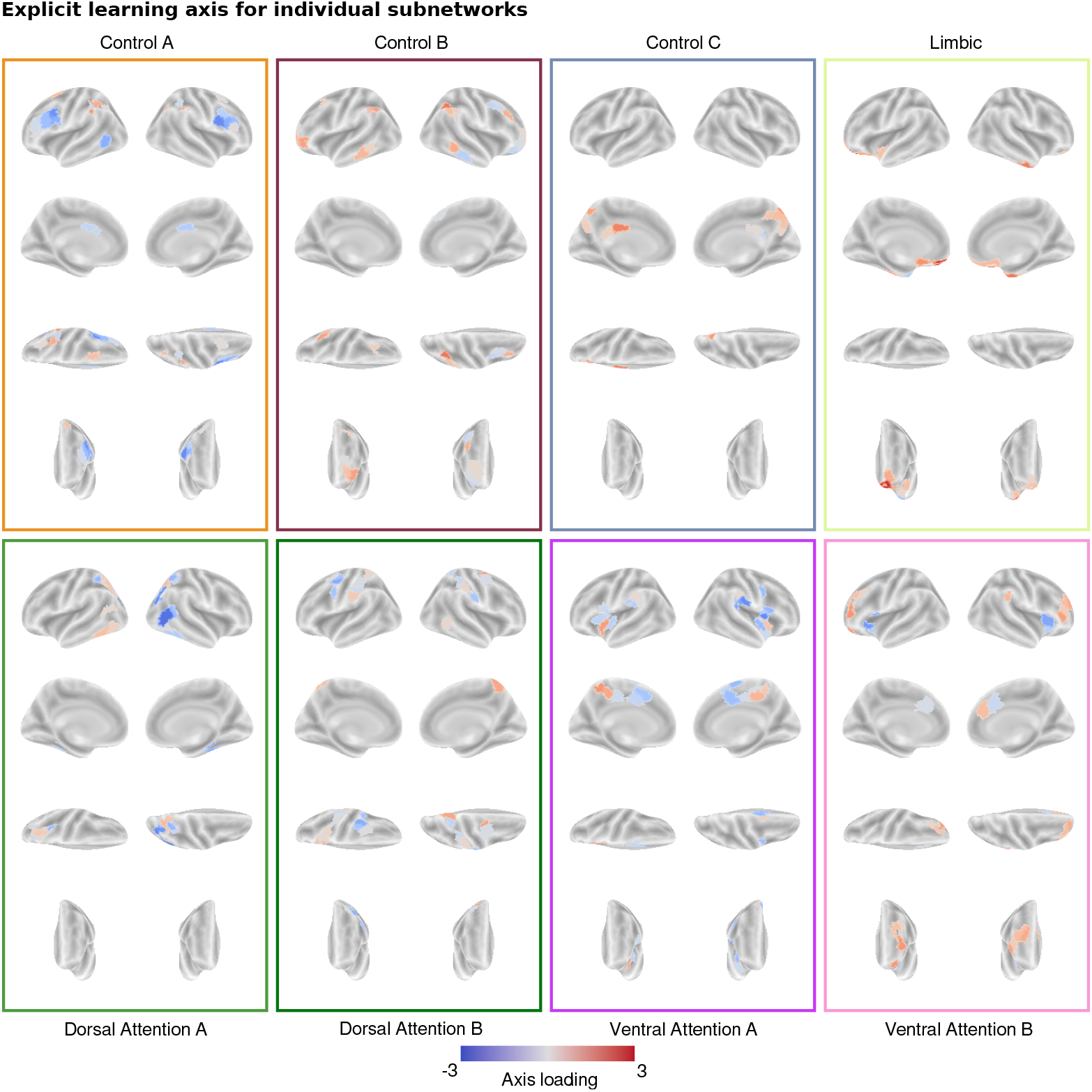
Explicit learning axis for individual subnetworks.

**Supplemental Figure 4:**
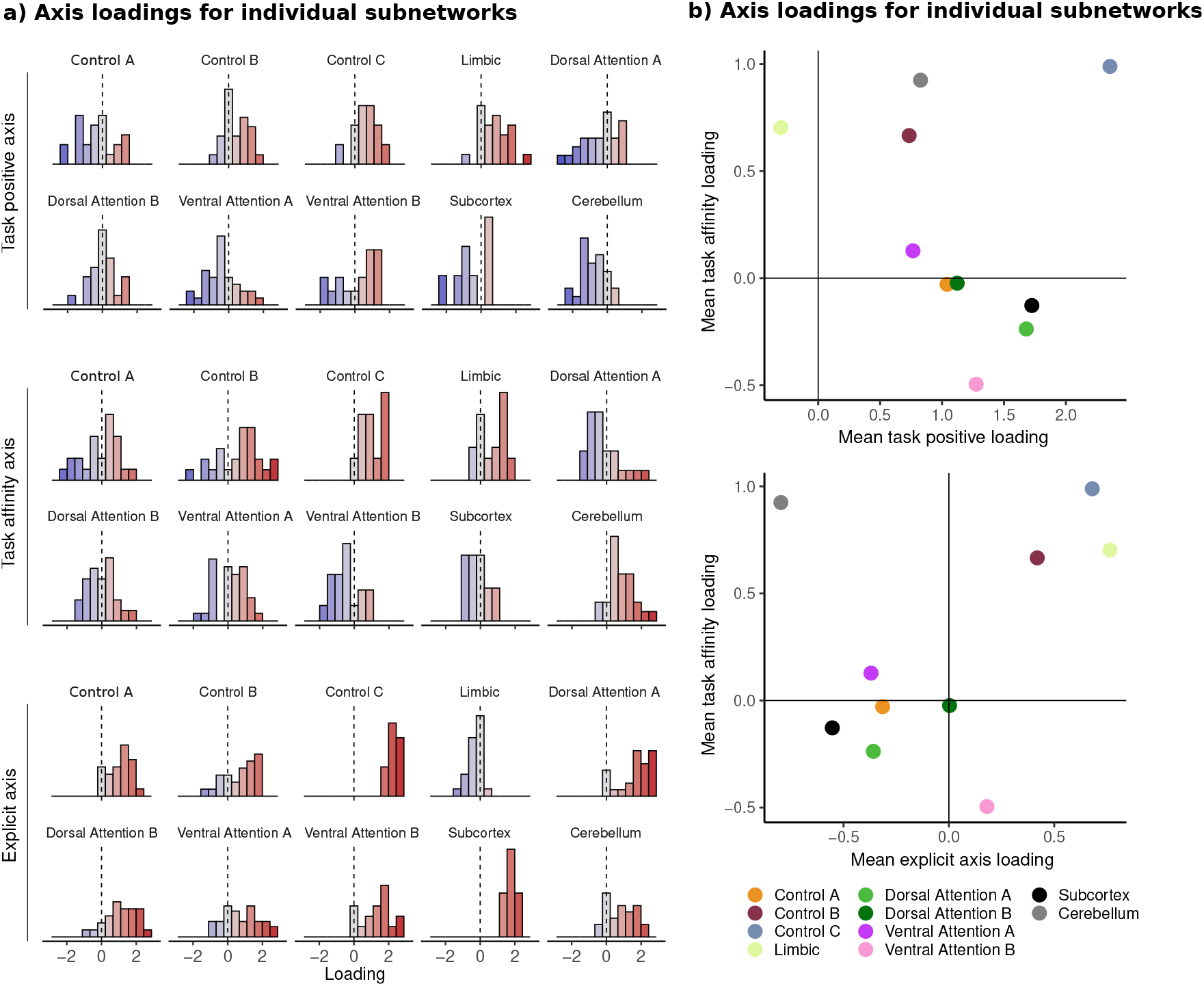
Loadings for the task positive, task affinity, and explicit learning axes - a) Loadings for individual ROIs in each subnetwork. b) Scatterplot of mean axis loadings for each subnetworks.

